# Temporally multiplexed imaging of dynamic signaling networks in living cells

**DOI:** 10.1101/2022.08.22.504781

**Authors:** Yong Qian, Orhan T. Celiker, Zeguan Wang, Burcu Guner-Ataman, Edward S. Boyden

## Abstract

Molecular signals interact to mediate diverse biological computations. Ideally one would be able to image many signals at once, in the same living cell, to reveal how they work together. Here we report temporally multiplexed imaging (TMI), which uses the clocklike properties of fluorescent proteins to enable different cellular signals to be represented by different temporal fluorescence codes. Using different photoswitchable fluorescent proteins to represent different cellular signals, we can linearly decompose a brief movie of the fluorescence fluctuations in a given cell, into a sum of the fluctuation traces of each individual fluorophore, each weighted by its respective signal amplitude. We demonstrate the power of TMI to report relationships amongst a diversity of second messenger, kinase, and cell cycle signals, using ordinary microscopes.

**One-Sentence Summary:** Imaging of many dynamic signals in a living cell is possible by using distinct clocklike fluorophores to represent the activity of each signal.

## Main text

Microscopy of living cells expressing fully genetically encoded fluorescent reporters of molecular signals is important for measuring the dynamics of these signals in relation to cellular states and functions. Multiplexed fluorescence imaging enables the important ability to see more than one signal at once, in a single living cell, so that relationships between the signals can be derived (*1–4*). Without this ability, it is hard to determine the relationships between different signals, key to understanding how they interact to yield cellular computations, and how such biological computations go wrong in disease states. As a simple example, suppose when signal A is high in a given cell, signal B goes low, and when signal A is low in a given cell, signal B goes high. Imaging of A and B in separate cells would miss out on this relationship; only by measuring them in the same cell will the relationship be easily seen. Of course, in most real biological signaling networks many more than two signals will interact to generate a biological computation. Traditionally, on the conventional microscopes commonly used in biology, multiplexed fluorescent imaging has relied on spectral differences between the fluorophores used to report different signals, which limits the number of signals observable to just a few. A recently developed multiplexing strategy uses self-assembling peptides to cluster dynamic fluorescent reporters at random, but stable, points throughout living cells, so they can be imaged separately by a conventional microscope (*1*). Such a spatially multiplexed imaging (SMI) strategy can measure cellular signals present at the specific locations of each cluster, with greater multiplexing than feasible with spectral multiplexing, raising the question of whether other kinds of dynamic signal, such as changes in cellular gene expression, changes in molecular location in a cell, and changes in the concentration of protein throughout a cell, might also be observable in a non-spectrally multiplexed, and thus potentially more scalable, way. From a design standpoint, the use of space as a resource to facilitate imaging in SMI – a theme used in other recent imaging developments, such as expansion microscopy (*5*) – raises the question of whether time could also be used as a resource to enhance the number of signals simultaneously observable in a single living cell, on a conventional microscope. Important for such an innovation to be impactful is ease of use in everyday biological contexts – for example, relying on fully genetically encoded fluorescent constructs, and on simple, widely available, conventional microscopes.

We here report that by using fluorophores that exhibit different clocklike temporal properties to report the expression of different genes, a brief movie of the summed fluorescence fluctuations can be acquired from a given cell, and then the individual gene expression levels can be isolated from the brief movie via standard linear unmixing algebra (**Fig. 1**). In the current study, we use reversibly photoswitchable fluorescent proteins (rsFPs) with different off-switching rates to achieve the clocklike effect, although in principle any fluorophore with temporal fluctuations could be adapted to this kind of concept. We show the utility of this method in analysis of the cell cycle, and furthermore demonstrate the use of this concept, which we call temporally multiplexed imaging (TMI), to measure many kinase activities at once. We show that TMI can support the imaging of many signals at once, more than feasible with traditional spectral multiplexing, revealing relationships between signaling proteins involved in the cell cycle that were not known before, but can now be easily mapped. TMI requires no hardware beyond standard epifluorescent or confocal microscopes commonly available to biologists, and can image many signals at once using a single-color channel, since the information is encoded in time, not through the spectrum.

**Fig. 1.**
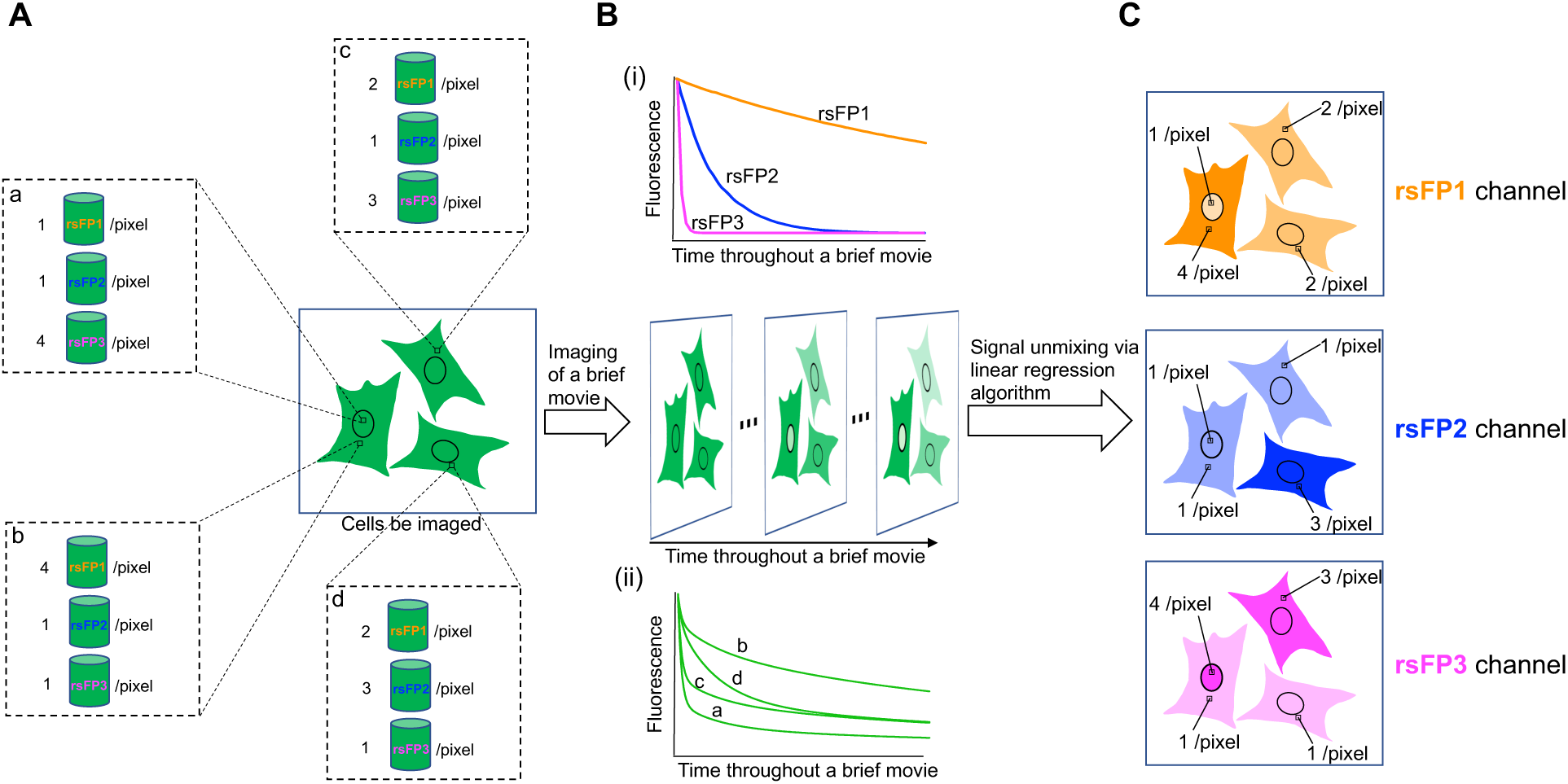
Concept of temporally multiplexed imaging (TMI), and implementation using reversibly photoswitchable fluorescent proteins (rsFPs). (**A**) Multiple rsFPs (denoted 1 through 3), even of the same color (note well, the colors of the text do not indicate the colors of the fluorophores, but are simply for reference by traces and schematics later in this figure), are expressed in cells. A cellular signal is indicated by the amount of a given fluorophore in a given cell (or part of a cell), e.g. because the fluorophore is expressed under a specific promoter, or fused to a specific protein, or translocated to a certain point. “a” through “d” represent different pixels exhibiting the indicated numbers of rsFP1, rsFP2, and rsFP3 molecules (“a” is in the nucleus of a cell; “b”, “c”, and “d” are in the cytosol). These rsFPs are spectrally indistinguishable by conventional microscopes. (**B**) Despite their spectral similarity, these rsFPs behave differently over time during continuous imaging (i) – in this study, due to differences in their photoswitching off-kinetics during continuous illumination, although other clocklike properties of fluorophores could be explored in the future. The reference traces of rsFP1 to rsFP3 are obtained individually, in calibration experiments, using the same imaging conditions as used in a later experiment probing a biological question. The fluorescence trace of each pixel acquired during a brief movie obtained during the experiment probing the biological question, then, would be a linear combination of all the rsFP traces summed at that pixel, each scaled by the number of fluorophores of that kind present at that pixel (ii). (**C**) The fluorescence intensity of each rsFP at each pixel is obtained via standard linear unmixing of the fluorescence trace acquired during the brief movie obtained during the experiment probing the biological question, using the reference traces acquired in the calibration experiments. The resultant images can then be pseudo-colored for easy visualization, by giving each rsFP a different digital color in software (here, orange, blue, and pink for rsFP1, rsFP2, and rsFP3, respectively).

In our initial implementation of TMI, multiple rsFPs, even of the same color, but exhibiting different clocklike or temporal behaviors (i.e., different off-switching rates during continuous imaging) are expressed in the same cell, for example under different promoters if dynamic gene expression monitoring is desired (**Fig. 1A**). Since each fluorophore behaves differently over time during continuous imaging (**Fig. 1Bi**), different pixels, which contain different concentrations of each fluorophore, will exhibit different overall timecourses of fluorescence over the timescale of a brief movie being recorded (**Fig. 1Bii**). By computationally unmixing the overall fluorescence trace at each pixel (**Fig. 1Bii**) into a linear weighted combination of the reference traces exhibited by each fluorophore alone (**Fig. 1Bi**; note well, these reference traces are derived in a separate experiment, with just one fluorophore present, but under the same imaging conditions), one can easily reconstruct the amplitude of each fluorophore at a given pixel – it is simply the weight associated with that fluorophore’s reference trace in the overall summed trace (**Fig. 1C**).

To validate this concept, we first explored green rsFPs, such as those whose fluorescence can be switched “off” by blue/cyan light and switched “on” by purple light (**Fig. 2A**). We first chose a set of green rsFPs with distinct off-switching kinetics, so that they could be distinguished in TMI imaging of cells (**Fig. 2B**). We found out that Dronpa (*6*), rsFastLime (*7*), and rsGreenF-Enhancer (rsGreenF-E) (*8*) exhibited highly different off-switching kinetics (**Fig. 2B**). Skylan-NS (*9*) had kinetics similar to rsFastLime (**Fig. S1A**), so we performed directed evolution (**Fig. S1A, B**) to identify a variant that we named Skylan62A (Skylan-NS-L62A), that exhibited photoswitching kinetics between those of rsFastLime and Dronpa (**Fig. S1A**), without changing spectral characteristics (**Fig. S1C**). We identified rsEGFP2-Enhancer (rsEGFP2-E) (*8, 10*) as an rsFP with off-switching kinetics between those of rsGreen-F-E and rsFastLime (**Fig. S1D**). Finally, we added a non-switching fluorescent protein, YFP (*11*), to create a 6th temporally distinguishable candidate (**Fig. 2B**).

**Fig. 2.**
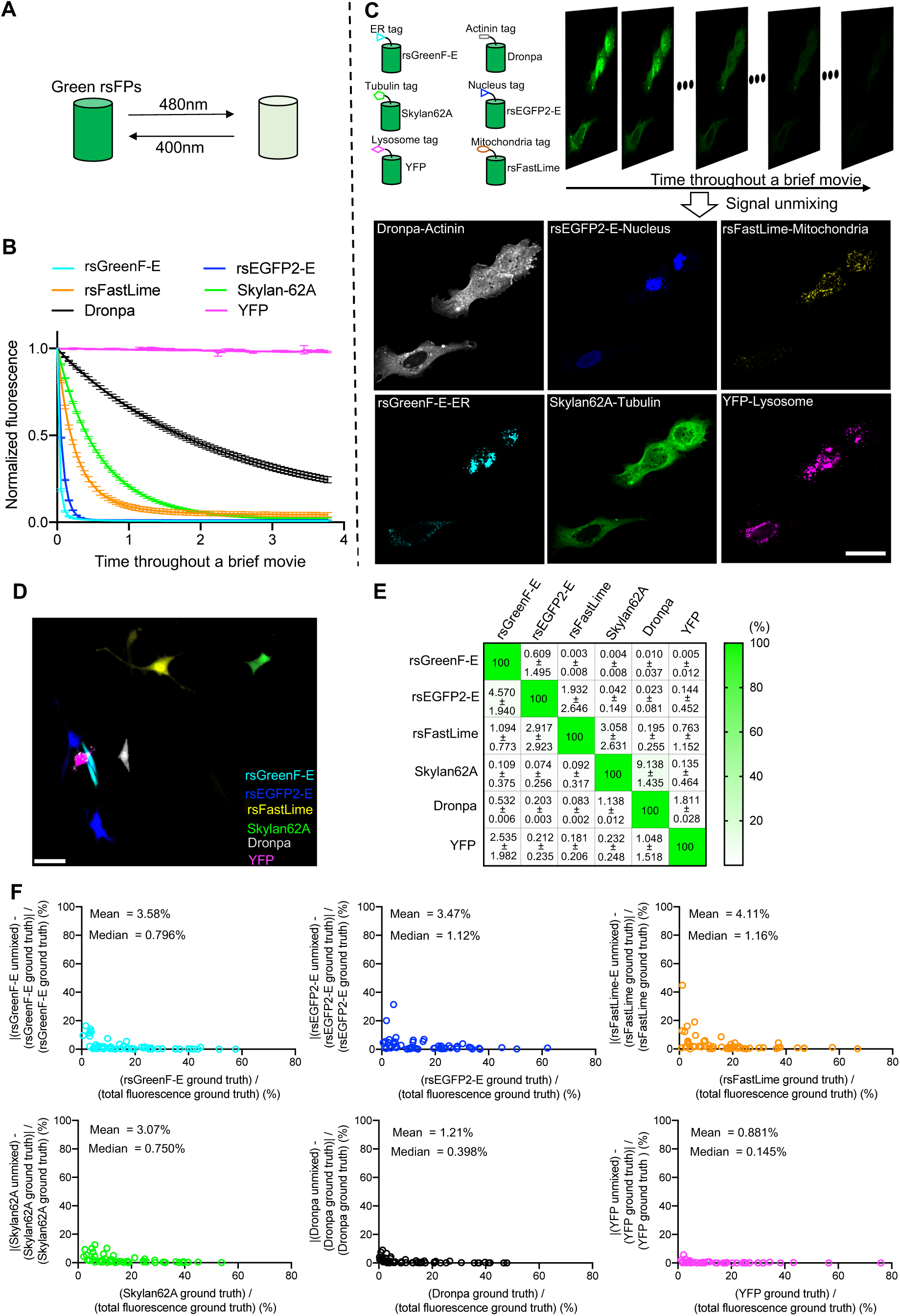
Temporally multiplexed imaging of six green fluorescent proteins (FPs). (**A**) Green rsFPs can be switched to “off” states during imaging with continuous excitation at ~480 nm and switched to “on” states by a pulse of purple light (~400 nm). (**B**) Reference traces of rsFastGreen-E, rsEGFP2-E, rsFastLime, Skylan-62A, Dronpa, and YFP in U2OS cells (cells were fixed, in this figure, for subsequent immunostaining, although typically such traces would be acquired in the living state), n=10-15 cells from 3 brief movies from 1 cell culture batch. Data are shown as mean ± standard deviation (SD). Illumination: 488 nm at 40 mW/mm^2^. (**C**) Co-expression of ER-targeted rsGreenF-E, nucleus-targeted rsEGFP-E, mitochondria-targeted rsFastLime, tubulin-targeted Skylan-62A, actinin-targeted Dronpa, and lysosome-targeted YFP in U2OS cells. A 70-frame brief movie of fixed U2OS cells was taken over 3.8 seconds, using the same imaging conditions and microscope as in **B**, and six images were obtained via linearly unmixing the fluorescence trace at each pixel in the brief movie, using the reference traces in **B.** Scale bar, 20 µm. (**D**) Representative merged image (out of 15 images taken from 2 cell culture batches) of live NIH/3T3 cells expressing all 6 green FPs of **B** (in this case, not fused to targeting tags) from six images unmixed from the source brief movie. Each cell was transfected with exactly one FP (see Methods for details). Scale bar, 50 µm. (**E**) FP-FP crosstalk, calculated from the linear decomposition coefficients extracted from 15 images from 2 cultures experimented upon as in **D**, and expressed as a percentage of the true FP brightness for a given cell (recall, each cell expresses exactly one FP, providing ground truth). Values are shown as mean ± SD; color represents mean. (**F**) Accuracy of linear decomposition of temporally multiplexed brief movies into individual FP images was simulated by taking actual cell images, and populating the pixels with traces that are randomly scaled versins of the curves of panel **B**, followed by unmixing and comparing to the ground truth (which, being simulated, is exactly known). See **Fig. S3** and Methods for details. Each data point represents one cell. The X-axis value of each data point represents the ground-truth fluorescence of an FP divided by the total ground-truth fluorescence summed over all FPs, for that cell. The Y-axis value of each data point represents the fluorescence difference between the unmixed image and its corresponding ground truth, divided by the ground truth.

We expressed the six aforementioned rsFPs in U2OS cells, each fused to a distinct, well-validated, subcellular targeting sequence so that each fluorophore would be targeted to a different biological structure. We fixed the cells (for subsequent immunostaining-based validation) and acquired brief (70-frame) movies over a short period, 3.8 seconds, on a standard confocal microscope. We then unmixed the brief movies using the reference traces (**Fig. 2B**, acquired under identical conditions, including fixation) using standard unmixing linear algebra (see **Methods**) and found that indeed we could identify the different structures, recapitulating known cellular morphologies (**Fig. 2C**). As a first strategy for validating the unmixing, we compared such resulting images to those obtained by antibody staining against a FLAG-tag fused to each of the six FPs in turn, and found no difference in the correlation between the rsFPs and the antibody staining when the rsFPs were expressed individually (which serves as a ground truth), vs. when they were expressed all at the same time (**Fig. S2A, B**). Thus, crosstalk between different rsFPs is minimal when analyzed in a temporally multiplexed way. As a second measure of TMI crosstalk, we expressed the chosen rsFPs in individual NIH/3T3 cells, with each cell expressing only one FP, to serve as a ground truth, and then measured the crosstalk of each FP to the other five FPs (**Fig. 2D**), finding crosstalk to be in the few percent range, and often much lower (**Fig. 2E**). Finally, as a third, independent, validation, we simulated a 70-frame video based on a fluorescence image of NIH/3T3 cells, with each pixel exhibiting a simulated linear combination of the six fluorophores of **Fig. 2B** (**Fig. S3A**), thus serving as a ground truth. We then simulated a brief movie of the fluorescence dynamics, including added noise, then unmixed the brief movie as above to obtain the reconstructed single-fluorophore images (**Fig. S3B**). We obtained near-perfect Pearson’s correlation values between unmixed images and ground truth images – exceeding 99.8% (**Fig. S3B**). Analysis of simulated reconstructions of individual cell data (**Fig. 2F**) showed that most cells had deviations from the ground truth of a few percent (although for extremely dim cells, the error, as expected, could be greater). Thus, in several independent validations, we found that linear unmixing of summed brief movies of cells containing mixtures of fluctuating fluorophores yielded highly accurate reconstructions of signals, when compared to ground truth datasets or simulations.

We next explored the use of off-switching red rsFPs for TMI. Off-switching red rsFPs are a type of FPs whose fluorescence can be switched to a dim state when illuminated with yellow/orange light and will resume bright fluorescence under illumination by blue/cyan or purple light (**Fig. 3A**). We selected rsTagRFP (*12*) and rsFusionRed1 (*13*) as two initial candidates for TMI, based on their switching rates (**Fig. 3B**). We engineered a new rsFP, which we named rScarlet, from mScarlet (*14*), inspired by the development of rsCherryRev1.4 from mCherry (*15, 16*) (**Fig. S4A**). rScarlet exhibited slower photoswitching kinetics than rsTagRFP and rsFusionRed1, and similar spectral properties to mScarlet (**Fig. S4B, C, Fig. 3B**). Next, we selected a non-switching RFP, mCherry (*17*), as the fourth fluorophore for TMI. Many other commonly used RFPs exhibited substantial off-photoswitching with excitation by yellow/orange light, and may be useful in the future (**Fig. S5**). We expressed the four chosen red FPs in U2OS cells, with each FP targeting a different subcellular structure, fixed the cells, and acquired brief (70-frame) movies over 8.6 seconds, on a standard confocal microscope. After linear unmixing, each unmixed image showed the expected subcellular structure (**Fig. 3C**), with minimal crosstalk when validated with antibody staining against epitope-tagged indicators (**Fig. S6**). When the four FPs were expressed in NIH/3T3 cells individually, the crosstalk between each pair of FPs was in the few percent range or lower (**Fig. 3D, E**). Repeating the simulation described above revealed that, as with green rsFPs, red rsFPs could yield extremely low crosstalk (**Fig. 3F, Fig. S7**). Thus, as with green rsFPs, red rsFPs could support well-separable TMI signals obtained with low crosstalk, when validated in multiple, independent ways.

**Fig. 3.**
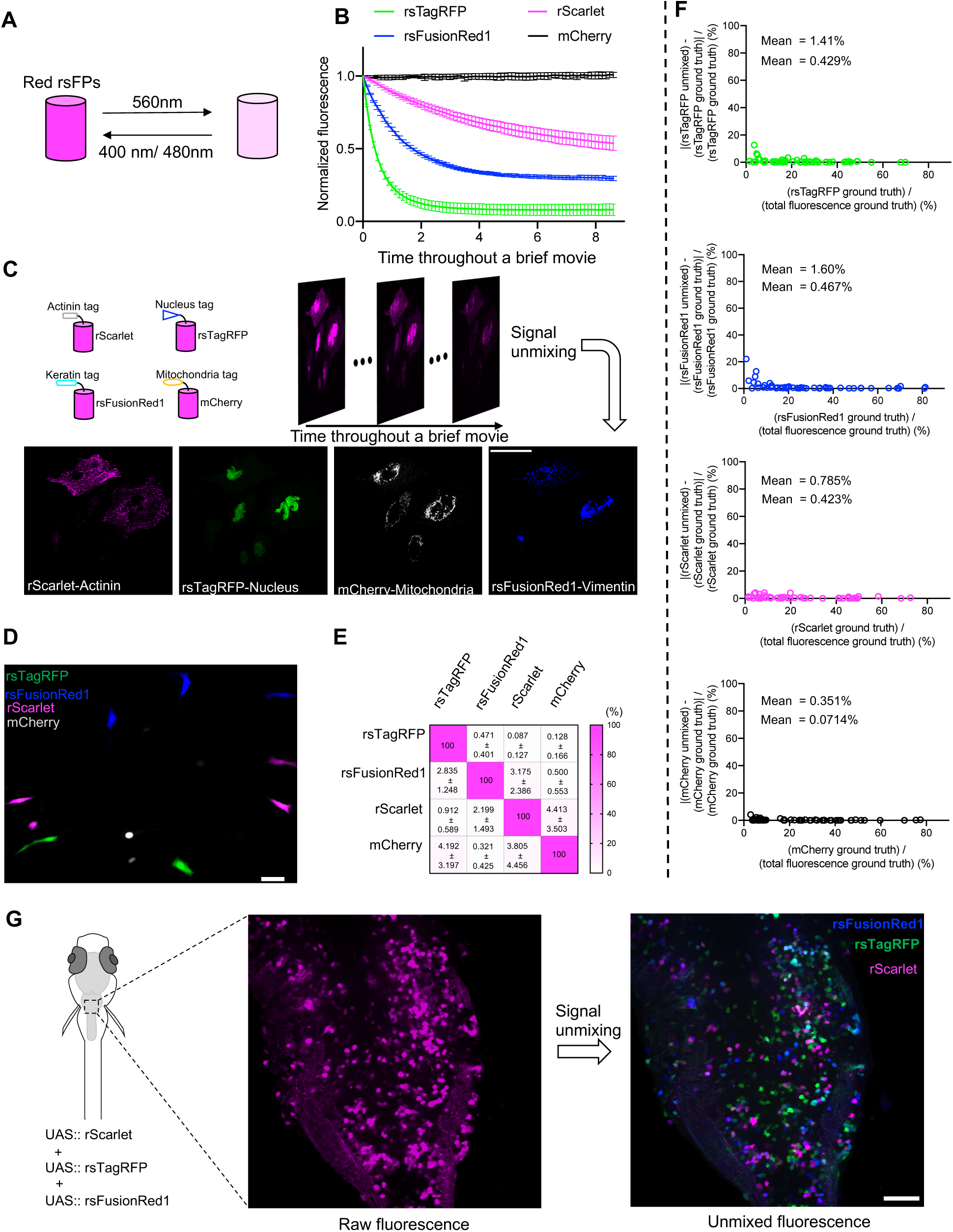
Temporally multiplexed imaging of red FPs. (**A**) Red rsFPs switch “off” with excitation at ~560 nm and switch “on” with irradiation with either blue (480 nm) or purple light (400 nm). (**B**) Reference traces of rsTagRFP, rsFusionRed1, rScarlet, and mCherry in U2OS cells (cells were fixed for subsequent immunostaining as in **Fig.2B**). Traces were collected from 10-15 cells from 1 cell culture batch, and are shown as mean ± SD. Illumination: 561 nm at 50 mW/mm^2^. (**C**) Co-expression of nucleus-targeted rsTagRFP, vimentin-targeted rsFusionRed1, actinin-targeted rScarlet, and mitochondria-targeted mCherry in U2OS cells. A 70-frame brief movie of fixed U2OS cells was taken in 8.6 seconds, and images of the four FPs were obtained via linearly unmixing the fluorescence trace at each pixel in that brief movie using the reference traces in **B.** Scale bar, 20 µm. (**D**) Representative merged image (out of 6 images taken from 2 cell culture batches) of live NIH/3T3 cells with expression of 4 red FPs (not fused to targeting tags) from four images unmixed from the source brief movie. Each cell was transfected with exactly one FP (see Methods for details). Scale bar, 50 µm. (**E**) FP-FP crosstalk was calculated from the linear decomposition coefficients extracted from 6 images from 2 cultures experimented upon as in **D**, and expressed as a percentage of the true FP brightness for a given cell (recall each cell has exactly one FP). Values are shown as mean ± SD; color represents the mean. (**F**) Accuracy of linear decomposition of temporally multiplexed movies into individual FP images was simulated by taking cell images and populating the pixels with traces that are scaled versions of the curves of panel **B**, followed by unmixing and comparing to the ground truth – which, being simulated, are exactly known. See **Fig. S7** and Methods for details. Each data point represents one cell. The X-axis value of each data point represents the ground-truth fluorescence of a FP divided by the total ground-truth fluorescence, for that cell. The Y-axis value of each data point represents the fluorescence difference between the unmixed image and its ground truth, divided by the ground truth. (**G**) Single-color “brainbow” in the larval zebrafish brain, based on temporally multiplexed imaging of rScarlet, rsTagRFP and rsFusionRed1. Middle, representative raw fluorescence image (out of 20 images taken from 3 animals) of zebrafish larvae hindbrain (dorsal view) at day 5 post fertilization; right, brainbow-like image obtained via linear decomposition of the brief movie acquired, showing the same area as the image in the middle panel. Scale bar, 20 µm.

Brainbow, the combinatorial expression of fluorescent proteins in neurons for neural identification and tracing (*18*), is popular, but requires multispectral imaging. We asked whether TMI could support a “single color brainbow” strategy, enabling one microscope color channel to identify many different cells expressing different combinations of fluorophores. We transiently expressed three red fluorophores, rsTagRFP, rsFusionRed1, and rScarlet in the nervous system of zebrafish larvae, in a mosaic fashion (*19, 20*). By imaging all three fluorophores over the period of a brief movie, in a single color channel, and then unmixing the fluorophores into separate channels, we could obtain brainbow-like images in living zebrafish brain via TMI (**Fig. 3G, Fig. S8**). In contrast to standard brainbow technology, which requires many color channels to be used, TMI-based brainbow only requires one color channel, and thus could free up other optical channels for synergistic purposes such as imaging of Ca^2+^, voltage, or neurotransmitters, or optogenetics, things which are difficult to do if many color channels have been already reserved for brainbow anatomy.

We next explored whether TMI could help with the visualization of many dynamical signals in living cells. Fluorescent, ubiquitination-based cell cycle indicator 4 (FUCCI4) is a previously published indicator system that reports all four cell cycle phases using cell cycle-regulated proteins fused to spectrally distinct FPs (*21*). If a TMI version of FUCCI4 were possible, then the freed-up spectrum could be used for other fluorescent reporters, perhaps to examine how other signaling cascades influence the cell cycle. Here we developed a single-color version of FUCCI4 by replacing the four FPs in the original FUCCI4 with four FPs that we characterized above for TMI (**Fig. 4A**). Specifically, Dronpa, YFP, rsGreenF-E, and Skylan62A were fused to the cell cycle-regulated proteins Cdt_130-120_, SLBP_18-126_, Geminin_1-110_, and histone H1.0 respectively. Analogous to the original FUCCI4, the G1-S transition is reported by the emergence of rsGreenF-E fluorescence while YFP fluorescence persists, and the S-G2 transition is marked by the loss of YFP amidst stable rsGreenF-E fluorescence. Chromosome condensation, as reported by Skylan62A, indicates the M phase; finally, loss of rsGreenF-E fluorescence and the appearance of Dronpa and YFP fluorescence means the beginning of the G1 phase. We imaged cell cycle transitions in NIH/3T3 cells using our single-color FUCCI4 system and were able to identify each transition in the cell cycle just as with the original FUCCI4 (**Fig. 4B, C, Fig. S9**).

**Fig. 4.**
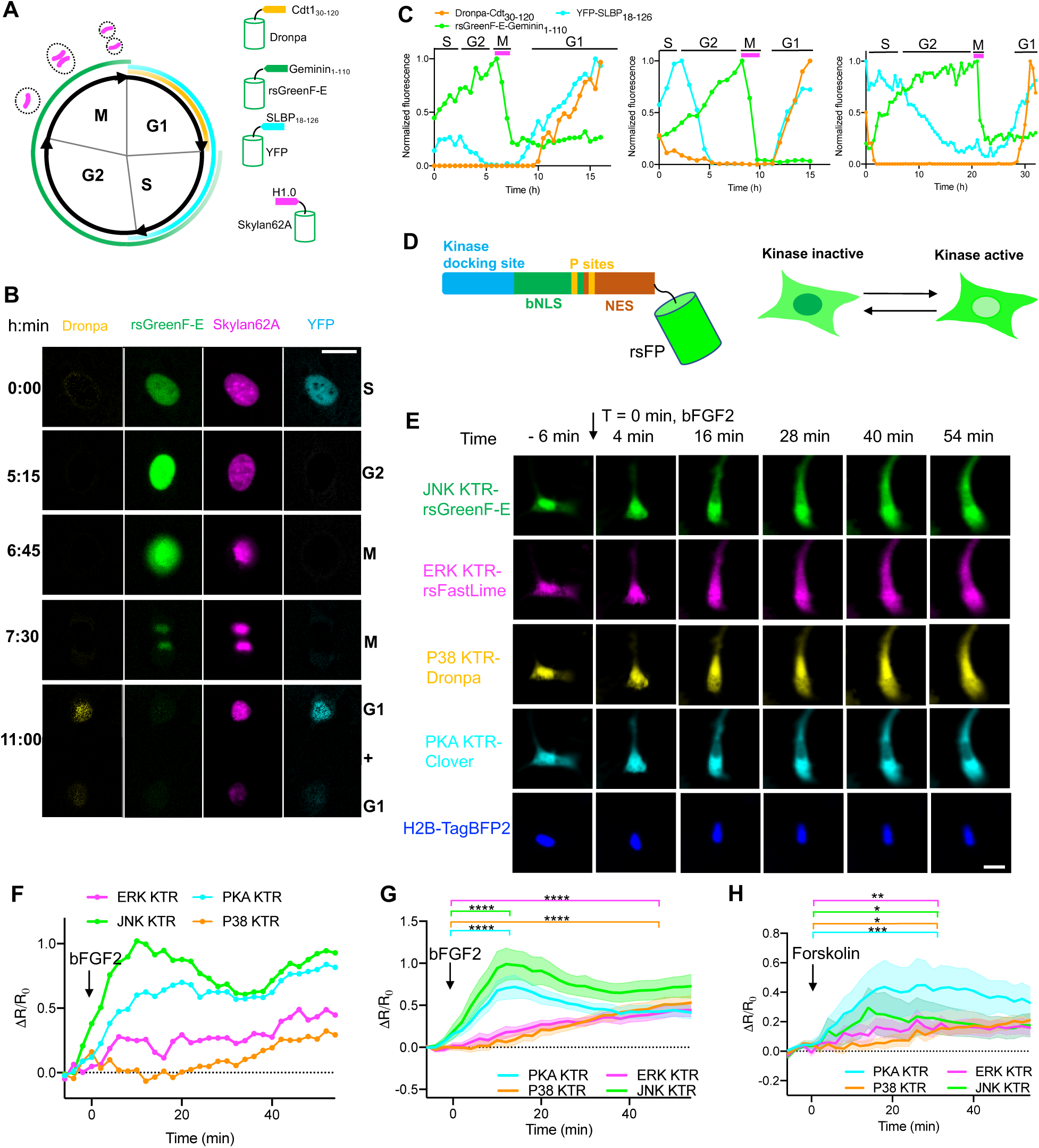
Temporally multiplexed imaging allows simultaneous observation of many biological signals at once in a single living cell. (**A**) Diagram of cell cycle indicators (known as fluorescent, ubiquitination-based cell cycle indicator 4, abbreviated FUCCI4). Dronpa, YFP, rsGreenF-E and Skylan62A were fused to cell cycle-regulated proteins Cdt1_30-120_, SLBP_18-126_, Geminin_1-110_, and histone H1.0, respectively. Analogous to the original FUCCI4, the G1-S transition is reported by the emergence of rsGreenF-E fluorescence while YFP fluorescence persists, and the S-G2 transition is marked by the fluorescence loss of YFP and stable rsGreenF-E fluorescence. Chromosome condensation, labeled by Skylan62A, indicates the M phase; finally, loss of rsGreenF-E fluorescence and the appearance of Dronpa and YFP fluorescence means the beginning of the G1 phase. (**B**) Tracking of cell cycle phase in NIH/3T3 cells via time-lapse imaging (i.e., taking brief movies at different points in time, each of which becomes a distinct datapoint in the time lapse) using TMI-based single-color FUCCI4. A mother cell dividing into two daughter cells was captured during a 11-hour imaging session, with brief movies acquired at each of the indicated times. Scale bar, 20 µm. (**C**) Fluorescence traces of Dronpa-Cdt1_30-120_ (orange), rsGreenF-E-Geminin_1-110_ (green), and YFP-SLBP_18-126_ (Cyan) of 3 cells over their cell divisions, tracing one arbitrary daughter cell from each mother cell after M phase. Fluorescence was normalized to maximum value over the time-lapse period. Magenta bars indicate time of observation of chromosome condensation. Cell-cycle phases were assigned as depicted in **A**. (**D**) Diagram of TMI-based kinase translocation reporters (KTRs). A KTR contains a kinase docking site, a phospho-inhibited bipartite nuclear localization signal (bNLS) containing phosphorylation sites (P sites), a phospho-enhanced nuclear export signal (NES) containing P sites, and an rsFP. When the corresponding kinase is inactive, the KTR protein is unphosphorylated and is nuclear enriched. When the corresponding kinase is active, the KTR protein is phosphorylated and excluded from the nucleus. (**E**) Four kinase sensors were made using four green FPs: an rsFastLime-based ERK sensor, an rsGreenF-E-based JNK sensor, a Dronpa-based P38 sensor, and a Clover-based PKA sensor. NIH/3T3 cells expressing all four KTRs and H2B-TagBFP as a nuclear marker were imaged and stimulated with mouse basic fibroblast growth factor 2 (bFGF2, 20 ng/ml), which is known to drive JNK, P38, ERK, and PKA. A representative cell (out of 16 cells taken from two cell culture batches) is shown (four pseudo-channels unmixed from the green channel, and one blue channel) at the indicated time points. Scale bar, 20 µm. (**F**) Activity traces of the four kinases from the representative cell in **E**. R, ratio of cytoplasmic intensity to nuclear intensity. Change in fluorescence of the sensors is plotted as ΔR/R_0_ (R_0_ was the averaged value from t = −6 min to t= −4 min). (**G**) Averaged traces of four kinase activities recorded from NIH/3T3 cells, with 20 ng/ml bFGF2 stimulation at t = 0 min; n = 16 cells from two culture batches. Data are shown as mean ± standard error of the mean (SEM). Wilcoxon rank sum tests were run between the averaged values from t = −6 min to t = 0 min and the averaged values from t = 12 min to t = 18 min for PKA KTR and JNK KTR. Wilcoxon rank sum tests were run between the averaged values from t = −6 min to t = 0 min and the averaged values from t = 48 min to t = 54 min for ERK KTR and P38 KTR. Traces of individual cells are shown in **Fig. S10**. (**H**) Averaged traces of four kinase activities recorded from NIH/3T3 cells, with stimulation with 50 µM forskolin at t = 0 min; n = 10 cells from two culture batches. Data are shown as mean ± SEM. Wilcoxon rank sum tests were run between the averaged values from t = −6 min to t = 0 min and t = 30 min to t = 36 min for all KTRs. Traces of individual cells are shown in **Fig. S11**. Throughout the figure: *p< 0.05; **p < 0.01; ***p < 0.001, ****p < 0.0001.

We next explored whether TMI could help with the imaging of signals other than gene expression, such as signals reported by the movement of a target protein from one cellular location to another. Kinase translocation reporters (KTRs) are fluorescent sensors that report protein phosphorylation by translocating between the cytoplasm and the nucleus (*22*) (**Fig. 4D,** left). When a kinase of interest is inactive, the KTR for that kinase is unphosphorylated, which leads to its nuclear localization; when the kinase of interest is active, phosphorylation of the KTR leads to translocation of the reporter to the cytoplasm (**Fig. 4D,** right). We replaced the FPs in the original KTRs for JNK, ERK, and P38, with rsGreenF-E, rsFastLime, and Dronpa respectively. Together with PKA KTR-Clover (Clover being a non-photoswitching FP), we performed TMI with four kinase sensors simultaneously. NIH/3T3 cells expressing all four KTRs and H2B-TagBFP as a nuclear marker were imaged and stimulated with mouse basic fibroblast growth factor 2 (bFGF2, 20 ng/ml), which is known to drive kinases such as JNK, P38, ERK, and PKA (*23–25*). We observed a fast onset of JNK activity, as well as of PKA activity, both of which tapered off slightly over time, whereas ERK and P38 activities steadily increased over time (**Fig. 4E-G, Fig. S10**). In contrast, forskolin preferentially induced PKA activity, inducing the other three kinases to a lesser degree, as compared to the bFGF2 case (**Fig. 4H, Fig. S11**), consistent with what has been observed previously (*2, 27, 28*). Thus, TMI allows many kinases to be imaged at once, facilitating comparison of their trajectories in individual cells, and examination of the temporal relationships in their activities, important for understanding how these kinases work together in a signaling network. Future studies systematically mapping out kinase pathways with TMI could help with the investigation of many healthy processes and disease states.

Given that one easily deployable impact of TMI is that it enables sets of existing reporters of expressed genes (**Fig. 3G, Fig. 4A-C**) or dynamical signals (**Fig. 4D-H**), easily converted to TMI form just by swapping out the old fluorophores with TMI replacements, to be simultaneously measured using a single color channel, an immediate next step is to see whether the newly freed up spectral bandwidth could be used to incorporate additional reporters, to measure more relationships than previously possible. We first combined our single-color TMI version of FUCCI4 with conventional reporters of cyclin-dependant kinases 2 (CDK2, based on TagBFP2) (*29*) and 4/6 (CDK4/6, based on mCherry) (*30*) (**Fig. 5A**, left). As with the kinase imaging we reported earlier, these two CDK reporters stay in the cell nucleus when their respective kinases are inactive, and translocate to the cell cytoplasm when activated. Cell cycle progression is cooperatively regulated by multiple CDKs, and thus knowing how CDK activities proceed throughout each cell cycle phase is critical to understanding the molecular computations of cell cycle regulation. Past results indicated that both CDK2 and CDK4/6 enter the M phase with a high level of activity, which drops rapidly during mitosis until the cell reaches the early G1 phase, followed by a slow buildup of activity throughout the G1 phase (**Fig. 5A**, right) (*29–32*). Although it was known that both CDK2 and CDK4/6 activities are higher at the end of the G2 phase than at the beginning of the S phase (*27, 28*), how the amplitudes of CDK2 and CDK4/6 activities change throughout the timecourses of S and G2 were unknown. Here, enabled by TMI-based FUCCI4, we were able to easily observe how CDK2 and CDK4/6 activities change in all four cell cycle phases, including the S phase and G2 phase, in NIH/3T3 cells (**Fig. 5B, Fig. S12**). When we averaged the CDK signal amplitudes during three periods – the early, middle and late stages of each cell cycle phase – we replicated prior observations that CDK2 and CDK4/6 activity went through an abrupt drop in M phase, reaching a maximum dip in early G1, and then started to build up from a low level throughout G1 (**Fig. 5C, D**). In addition, we found that after entering S phase, CDK2 activity kept rising at a slow rate (**Fig. 5C, Fig. S13**)); the rise continued during G2, until reaching the peak observed in early M phase (**Fig. 5C, E**). However, CDK4/6 activity exhibited a different pattern. It plateaued throughout the S phase, never increasing beyond the initial peak in early S, only beginning to increase in amplitude when G2 began (**Fig. 5D, E**). These insights into the relationship between CDK2 and CDK4/6 activity in S and G2 phases may point to the ability to tease apart how they operate together for cell cycle regulation.

**Fig. 5.**
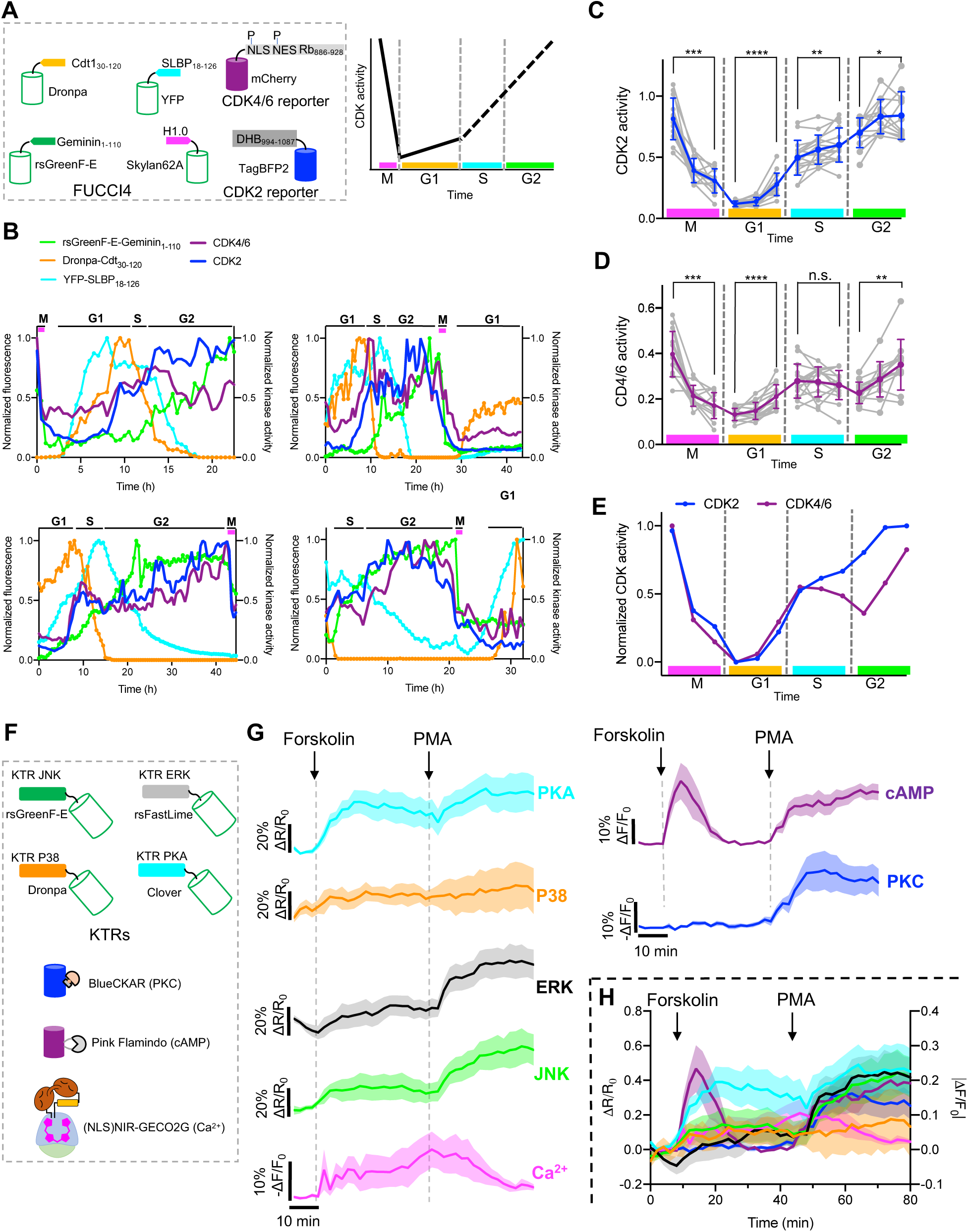
Combined temporal and spectral multiplexing for simultaneous imaging of large numbers of signals within single living cells. (**A**-**D**) Simultaneous observation of cell cycle phase changes and kinase activity. (**A**) Left, diagram of combined use of green rsFP-based FUCCI4 with a TagBFP2-based CDK2 reporter and an mCherry-based CDK4/6 reporter. Right, schematic of CDK activity in different cell cycle phases. CDK activity is higher at the end of G2 phase than that at the beginning of S phase, but the detailed CDK activity progression during S and G2 phases was unknown (dashed line) previously (*32*). (**B**) Representative traces of four fluorescence signals from FUCCI4, and two CDK signals from their respective reporters, over the cell cycle. Traces of four dividing NIH/3T3 cells are shown. Cells had brief movies acquired every 30 min, without stimulation. Fluorescence of Dronpa-Cdt1_30-120_ (orange), rsGreenF-E-Geminin_1-110_ (green), and YFP-SLBP_18-126_ (Cyan) was normalized to their maximum values. Cell cycle phases were determined by FUCCI4 signals using the principles described in **Fig. 4A**; magenta bars indicate observation of chromosome condensation. CDK2 activity (blue) was calculated as cytoplasm to nucleus fluorescence ratio; CDK4/6 activity (purple) was calculated as was CDK2 activity, followed by signal processing as in the previous paper (*30*). Both CDK2 and CDK4/6 traces were normalized to their maximum value for comparison. More traces from individual cells are shown in **Fig. S12**. (**C, D**) Plots of CDK2 (**C**) and CDK4/6 (**D**) activity as a function of stage (early, middle, and late stage) of each cell cycle phase. CDK2 and CDK4/6 activities were calculated as in **B,** without normalization. Each cell cycle phase was divided into early, middle and late stages evenly, and CDK activity for a given stage was obtained by averaging CDK values across that stage (see Methods). Data are shown as mean ± SD (blue, CDK2; purple, CDK4/6), with all individual values plotted as dots (grey; dots of the same cell cycle phase of the same cell are connected); n = 25 cells from 5 culture batches (note well, not all cells exhibited all complete phases). Wilcoxon signed-rank test with Holm-Bonferroni correction, ****p < 0.0001, ***p < 0.001, **p < 0.01, *p < 0.05, n.s., not significant. (**E**) Superimposed plots of CDK2 and CDK4/6 activity. Data points were from the mean values of the plots in **C** and **D**, and then normalized between 0 and 1 for visual comparison. (**F**-**H**) Simultaneous imaging of 7 signals within single NIH/3T3 cells. (**F**) Diagram of combined use of green rsFP-based KTRs with BlueCKAR, Pink Flamindo and NIR-GECO2G. (**G**) Signals of PKA (cyan), P38 (orange), ERK (black), JNK (green), Ca^2+^ (magenta), cAMP (purple), and PKC (blue) under stimulation with 50 µM forskolin (added at t = 8 min), and 100ng/ml PMA (added at t = 44 min); n = 7 cells from 2 culture batches, values are shown as mean ± SEM. Signals from individual cells are shown in **Fig. S14**. (**I**) Superimposed traces of **H,** for visual comparison.

As another example of how using TMI for a set of reporters could open up spectrum for additional signals to be measured, we combined our previously mentioned green FP-based KTRs with NIR-GECO2G (*33*), Pink Flamindo (*34*), and BlueCKAR (*2*), with the goal of simultaneously observing the following seven cellular signals within single cells: JNK, ERK, P38, PKA, Ca^2+^, cAMP, and PKC (**Fig. 5F**). NIH/3T3 cells expressing all seven reporters were imaged before and after the addition of 50 µM forskolin, followed by the addition of 100 ng/ml phorbol 12-myristate 13-acetate (PMA) 36 min later (without removal of the forskolin). Forskolin is known to induce substantial cAMP, Ca^2+^, and PKA responses, lesser ERK, P38, and JNK responses, and no PKC response (*1, 2*); the TMI imaging of all seven signals at once under forskolin challenge was consistent with these prior findings (**Fig. 5G, H, Fig. S14**). PMA is a commonly used PKC activator, but studies have shown that it could also activate PKA, cAMP, JNK, and ERK (*35, 36*). We found that, when PMA was added 36 minutes after forskolin challenge, PKC showed a rise, and PKA, cAMP, JNK and ERK showed boosted responses, whereas P38 and Ca^2+^ exhibited no change, and possibly a decline (**Fig. 5G, H, Fig. S14**). Thus, by combining spectral and temporal multiplexing we were able to image seven different signals in individual cells, to our knowledge more than previously possible with fully genetically encoded indicators being imaged in living cells on conventional fluorescent microscopes.

In summary, TMI in an easy-to-use, powerful, versatile, and inexpensive strategy for imaging potentially arbitrary numbers of signals in living cells, by taking advantage of the clocklike, temporal properties of fluorescent proteins. As shown here, TMI enables both functional and structural imaging of live specimens (and fixed specimens, if so desired) on standard epifluorescence or confocal microscopes comprising ordinary grade hardware. We were able to observe many signals at once, in individual cells, by using only one spectral channel via TMI, freeing up spectrum for imaging more signals. This ability enabled us to discover novel relationships between CDK2 and CDK4/6 in the cell cycle, and to observe seven different signals in single living cells.

As with any new technology, it is useful to think about how end users can use TMI in the most practical way. First, TMI relies on repeatable photoswitching of rsFPs, and thus even illumination of specimens is helpful for accurate signal unmixing (although, if uneven illumination cannot be avoided, in principle this could easily be calibrated out by measuring the reference trace on a pixel-by-pixel basis, amidst the uneven illumination, on control samples). Second, since the processing of reference traces and of final brief movies are done pixel-by-pixel, movement of imaging samples during each bout of photoswitching should be avoided; if movement occurs, image registration might be needed. Third, the imaging time for each brief movie, in the current study, ranged from 3 s to 16 s, depending on the illumination intensity; this sets the temporal precision of TMI in terms of measuring signals over time (e.g., TMI would not be currently useful for voltage imaging, which requires millisecond precision). In principle, stronger excitation could be used if a shorter time duration for the brief movie (such as < 1 s), and thus a faster “frame rate” for the overall measurement of signal over time, was preferred. Fourth, the brief movie must contain sufficient data for an accurate linear unmixing to be performed; based upon our current work, we recommend acquiring a 50 (or greater)-frame movie for each datapoint to be extracted. During such brief movies, with the illumination intensities utilized, the intensity of Dronpa dropped to 35% (or less) of its original fluorescence at the end of the brief movie utilized for green FP multiplexing (other green rsFPs typically had dropped essentially to 0% intensity by this point), and the intensity of rScarlet dropped to 60% (or less) of its original fluorescence at the end of the brief movie utilized for multiplexing of red FPs. Fifth, even though rsFPs can resume brightly fluorescing, with minimal loss, after each round of off-switching (*6–8,12,13*), we do note that the photobleaching of most FPs in TMI was somewhat larger than that seen in conventional imaging (**Fig. S15**), due to the prolonged illumination needed to take many brief movies one after the other, to extract a time-lapse dataset from a given set of living cells.

Future endeavors could computationally optimize the signal unmixing algorithm for higher accuracy and/or faster computing, and improve the brightness of currently available rsFPs, most of which are dimmer than commonly used non-switching FPs (**Table S1**). Functional indicators of calcium and other cellular messengers could be developed based on rsFPs, to broaden the number of signals measurable with TMI. If a set of bright photoswitchable small molecule fluorescent dyes were adapted for TMI, perhaps TMI could serve roles in other technology areas, e.g. the multiplexed immunofluorescence imaging of cleared and/or expanded samples. In summary, we think TMI is not only an immediately practical technique for biological imaging, but can be adapted by the community in many directions.

## Acknowledgements

We thank Panagiotis Symvoulidis for help with imaging setup and Jordan Harrod, Yangning Lu, and Bobae An for initial concept testing and useful discussions. We also thank Chi Zhang for cell culture preparation.

## Author contributions

Y.Q. and E.S.B conceived the project, designed the experiments, interpreted the data and wrote the paper. Y.Q. developed Skylan62A and rScarlet. O.T.C., and Z.W. wrote the code for TMI signal unmixing and TMI simulations. Y.Q. performed experiments in cultured cells. Y.Q., Z.W. and B.G.-A performed experiments in zebrafish. Y.Q. performed data analysis. E.S.B. supervised the project.

## Data and materials availability

All data that support the findings of this study are available from the corresponding author upon reasonable request. All sequences of genes created will be submitted to Genbank, and all plasmids constructed in the work will be provided to Addgene for free distribution.

## Supplementary Materials

### Materials and Methods

#### Molecular cloning

Plasmids used in this study were constructed by either restriction cloning or In-Fusion assembly. Sanger sequencing was used to verify DNA sequences. The genes of Dronpa (*6*), YFP (*11*), and mCherry (*17*) were amplified from Addgene plasmids 57260, 1816, and 55148 respectively. The genes for rsFastLime (*7*), rsGreenF (*8*), Skylan-NS (*9*), rsEGFP2 (*10*), rsTagRFP (*12*), rsFusionRed1 (*13*), and GFP enhancer nanobody (*8*) were synthesized de novo by Integrated DNA Technologies based on the reported sequences. Site-directed mutagenesis libraries were generated using Quikchange site-directed mutagenesis (Agilent). For expression in bacteria, genes were cloned into pBAD-HisD vector. For ubiquitous expression in mammalian cells, genes were cloned into plasmids with one of the three promoters: CMV promoter, EF-1α promoter, CAG promoter. For expression in zebrafish, genes were cloned into the pTol2-10xUAS backbone (for Gal4-dependent expression) (*19*).

All synthetic DNA oligonucleotides used for cloning were purchased from either Integrated DNA Technologies or Quintarabio. PCR amplification was performed using CloneAmp HiFi PCR Premix (Takara Bio). Restriction endonucleases and T4 DNA ligase were purchased from New England BioLabs and used according to the manufacturer’s protocols. In-Fusion assembly master mix (Takaro Bio) was used following the manufacturer’s instructions for plasmid In-Fusion assembly. Small-scale isolation of plasmid DNA was performed with plasmid mini-prep kits (Takara Bio); large-scale DNA plasmid purification was done by Quintarabio. Plasmids of rScarlet variants were isolated and purified using 96-well plasmid miniprep kits (Bioland Scientific LLC). Stellar Competent cells (Takara Bio) were used for cloning, small-scale DNA plasmid purification, and protein purification; DH5α or NEB Stable Competent cells (New England Biolabs) were used for large-scale DNA plasmid purification.

#### Cell culture and transfection

HEK293FT cells (Thermo Fisher) were grown and maintained in Dulbecco’s modified Eagle’s medium (DMEM) (Gibco) supplemented with 10% heat-inactivated fetal bovine serum (Gibco), 2 mM GlutaMax (Thermo Fisher Scientific), and 1% penicillin-streptomycin (Gibco), at 37 °C and 5% CO_2_. Cells were seeded on 24-well glass-bottom plates (Cellvis) or 96-well plates (Cellvis) before transfection. Transfection of HEK293FT cells was performed when cells were 40 - 60% confluent with TransIT transfection reagent (Mirus Bio) according to the manufacturer’s instructions. Briefly, for a 24-well plate well, 500 ng of plasmid DNA was mixed with 1.5 μl of TransIT reagent in 50 μl opti-MEM (Gibco). After 30-min incubation, the DNA and transfection reagent mix were added to the cell culture medium dropwise. Imaging was then performed 24 hours post-transfection.

U2OS cells (ATCC) were grown and maintained in McCoy’s 5A medium (Gibco) supplemented with 10% heat-inactivated fetal bovine serum (Gibco) and 1% penicillin-streptomycin (Gibco), at 37 °C and 5% CO_2_. The protocols for seeding and transfection of U2OS cells were the same as those used for HEK293FT cells. For transfection of multiple constructs, plasmids were added to opti-MEM with a total amount of 500 ng and an equal ratio. The plasmid DNA was then fully mixed by vortexing before TransIT transfection reagent was added. Imaging was performed 48 hours post-transfection.

NIH/3T3 cells (ATCC) were grown and maintained in Dulbecco’s modified Eagle’s medium (DMEM) (Gibco) supplemented with 10% bovine calf serum (Millipore Sigma) at 37 °C with 5% CO_2_. NIH/3T3 cells were tested for mycoplasma contamination every 3 months. NIH/3T3 cells were seeded on 24-well glass-bottom plates and transfection was performed when they were 40 - 60% confluent using Lipofectamine 3000 (Thermo Fisher), following the manufacturer’s instructions. For transfection of multiple constructs, equal amounts of plasmid constructs were fully mixed in opti-MEM (Gibco) via vortexing before the transfection reagent was added. Imaging was performed 16 to 48 hours post-transfection.

#### Screening of new reversibly photoswitchable fluorescent proteins

To screen a green photoswitchable fluorescent protein with an off-switching rate between Dronpa and rsFastLime, eight variants carrying mutations at position no. 62 of Skylan were constructed and transiently expressed in HEK293FT cells individually. A 70-frame brief movie was then recorded for each variant using an epifluorescence inverted microscope (Eclipse Ti-E, Nikon) equipped with an Orca-Flash4.0 V2 sCMOS camera (Hamamatsu) and a SPECTRA X light engine (Lumencor). NIS-Elements Advanced Research (Nikon) was used for automated microscope and camera control. Cells were imaged with a 40× NA 1.15 water-immersion objective lens (Nikon) at room temperature (excitation: 475/28 nm at 15 mW/mm^2^, emission: 525/50 nm). Purple light (390/22 nm at 2 mW/mm^2^ for 100 ms) was applied right before taking brief movies. The off-switching traces of each variant were extracted from 8 to 10 cells. The cells were chosen so that they were evenly distributed over the field of views. Skylan62A was the winner of the screening. The off-switching traces of rsGreenF, rsGreenF with the enhancer nanobody, rsEGFP2, and rsEGFP2 with the enhancer nanobody were also obtained using similar imaging setups and analysis (excitation: 475/28 nm at 5 mW/mm^2^, emission: 525/50 nm).

For the screening of a new red photoswitchable fluorescent protein with a slow off-switching rate, six mutations borrowed from rsCherryRev1.4 (*15,16*) were introduced to mScarlet followed by site-directed saturation at position no.148 and no.162 of mScarlet (*14*) using the following primer: 5’atgggctggttcgcgNNCaccgagcagttgtaccccgaggacggcgtgctgaagggccttKSCaagatggccctgcgcctg-3’. The plasmids of 196 variants were then amplified, isolated, and expressed individually in HEK293FT cells. Brief movies (70 frames) were then recorded (excitation: 555/28 at 9.4 mW/mm^2^, emission: 630/75 nm; on-switching: 475/28 nm at 9.6 mW/ mm^2^ for 100 ms) for the variants with detectable fluorescence on the same wide-field microscope used for the screening of new green photoswitchable fluorescent proteins. The off-switching traces of each tested variant were then extracted from 8 to 10 cells that were evenly distributed over the field of views. The winner of the screening was named rScarlet.

The photoswitching behaviors of commonly used red fluorescent proteins mScarlet (*12*), mRuby2(*37*), mApple (*38*), mCherry, mKate2 (*39*), tdTomato (*17*), TagRFP (*40*), stagRFP (*41*), FusionRed (*42*) and far-red fluorescent proteins mNeptune(*43*), mCardinal(*44*), mMaroon (*21*) were measured using the same imaging setups and analysis as those for rScarlet screening. Three photoswitching cycles were measured for each FP; a pulse of blue light (475/28 nm at 9.6 mW/mm^2^ for 50 ms) was used to switch FPs from the “off” state to the “on” state before each cycle.

#### Protein purification and in vitro characterization

To purify each protein sample for characterization, single E. coli colonies expressing each protein were picked and cultured in 2 mL liquid lysogeny broth (LB) medium supplemented with 100 µg/mL ampicillin at 37 °C overnight. This 2-mL culture was then inoculated into a 500 ml liquid LB medium supplemented with 100 µg/mL ampicillin and 0.02% L-arabinose (wt/vol) and cultured at 28 °C for 24 h. After culture, bacteria were harvested by centrifugation. Protein purification was then performed using Capturem His-tagged purification maxiprep kit (Takara bio) following the manufacturer’s instructions. Purified proteins were subjected to buffer exchange to 1X TBS (pH = 7.4) with centrifugal concentrators (GE Healthcare Life Sciences).

Absorption, excitation, and emission spectra of purified Skylan62A and rScarlet were measured using a Tecan Spark microplate plate. Extinction coefficients of Skylan62A and rScarlet were determined by first measuring the absorption spectrum of Skylan62A or rScarlet in 1X TBS. The concentration of each protein was then determined by measuring the absorbance of alkaline-denatured protein and assuming ε = 44,000 M^−1^cm^−1^ at 446 nm (*45*). The extinction coefficient (ε) of the protein was calculated by dividing the peak absorbance maximum by the concentration of protein. Proteins were switched to the “on” state with the illumination of purple light before each measurement.

To determine fluorescence quantum yields of Skylan62A and rScarlet, Skylan-NS and mScarlet-I were used as standards respectively. Briefly, the concentration of SKylan62A (or rScarlet) in 1X TBS was adjusted such that absorbance at the excitation wavelength was between 0.1 and 0.2. A series of dilutions of each protein solution and standard, with absorbance values ranging from 0.005 to 0.02, was prepared. The fluorescence spectrum of each dilution of each standard and protein solution was recorded and the total fluorescence intensities were obtained by integration. FPs were switched to their “on” state with the illumination of purple light before each measurement. Absorbance versus integrated fluorescence intensity was plotted for each protein and each standard. Quantum yield (Φ) was calculated from the slopes (S) of each line using the equation: Φ_protein_ = Φ_standard_ × (S_protein_/S_standard_).

#### Temporal multiplexing of rsFPs in U2OS cells for subcellular labeling

##### Temporal multiplexing of green rsFPs

rsGreenF-E was fused with an ER-targeting sequence (MLLSVPLLLGLLGLAVA) on the N-terminus and an ER-retention signal sequence (KDEL) on the C-terminus (*46*); rsEGFP2-E was fused with histone H2B on the N-terminus; rsFastLime was fused with a mitochondria targeting sequence (MSVLTPLLLRGLTGSARRLPVPRAKIHSL, from Addgene plasmid 57287) on the N-terminus; Skylan62A was fused with Tubulin (from Addgene plasmid 57302) on the C-terminus; Dronpa was fused with α-actinin (from Addgene plasmid 57260) on the N-terminus, YFP was fused with LAMP1 on the N-terminus (Addgene plasmid 1816) (*47*). Six more constructs were built by adding a FLAG-tag (DYKDDDK) to the C-terminus of each FP in the previous six constructs.

The six constructs with FLAG-tag were expressed in U2OS cells individually. In parallel, one of the FLAG-tagged constructs was coexpressed with the other five non-FLAG constructs in U2OS cells. Cells with the expression of subcellular compartment-targeted FPs were then fixed with 4% paraformaldehyde (PFA) 48 hours after transfection followed by two washes with 1X PBS and one wash with 1X Phosphate-Buffered Saline (PBS) containing 100 mM glycine at room temperature. Cells were permeabilized with 0.1% Triton X-100 for 10 minutes and then blocked with MAXBlock Blocking medium (Active Motif) for 15 min, followed by three washes for 5 minutes each at room temperature in 1X PBS. Next, samples were incubated with rabbit anti-FLAG antibody (Invitrogen) in MAXStain Staining medium (Active Motif) for 1 hour at room temperature followed by three washes for 5 minutes each at room temperature in 1X PBS. Then, samples were incubated with Alexa647-labeled goat-anti-rabbit antibody (Abcam) in MAXStain Staining medium (Active Motif) for 1 hour at room temperature followed by three washes for 5 minutes each at room temperature in 1X PBS. Samples were then stored in 1X PBS and imaged on a Nikon Eclipse Ti inverted microscope equipped with a spinning disk confocal (CSU-W1), a 40×, 1.15 NA water-immersion objective, and a 5.5 Zyla camera (Andor), controlled by NIS-Elements AR software.

For the imaging of the cells with only one FLAG-tagged construct expressed, two snapshot images were taken from the green channel (exposure time 50ms, excitation: 488nm, emission: 525/30 nm) and far-red channel (exposure time 50ms, excitation: 637nm, emission: 700/50nm) for each field-of-view (FOV). A colocalization test was then run between the two images from the same FOV to get a Pearson’s correlation value. For the imaging of the cells with 6 constructs expressed (one FLAG-tagged construct plus five constructs without FLAG tag), a 70-frame brief movie and a snapshot image were taken from the green (exposure time 50ms, excitation: 488nm at 40 mW/mm^2^, emission: 525/30 nm) and far-red channel, respectively for each FOV. Six unmixed images were obtained via signal unmixing of each brief movie. The unmixed image from the FLAG-tagged construct was then colocalized with the image taken from the same FOV via far-red channel to get a Pearson’s correlation value. Purple light (405 nm at 9.7 mW/mm^2^) was applied for 50 ms before brief movies were taken.

##### Temporal multiplexing of red FPs

rsTagRFP was fused with histone H2B (same as the tag fused to rsEGFP2-E in the previous experiment) on the N-terminus, rsFusionRed was fused to Human Vimentin Sequence (from Addgene plasmid 57306) on the C-terminus. rScarlet was fused with α-actinin (same as the tag fused to Dronpa in the previous experiment) on the N-terminus. mCherry was fused with a mitochondrial targeting sequence (same as the tag fused to rsFastLime in the previous experiment) on the N-terminus. Four more constructs were built by adding a FLAG-tag (DYKDDDK) to the C-terminus of each FP in the aforementioned four constructs.

The protocols for cell transfection, fixation, and immunostaining were the same as those for green FPs. Imaging of red FPs was also performed on the same microscope as the imaging of the green FPs. For the imaging of the cells with only one FLAG-tagged construct expressed, two snapshot images were taken from the red channel (exposure time 100ms, excitation: 561nm, emission: 579/34 nm) and far-red channel (exposure time 50ms, excitation: 637nm, emission: 700/50nm) for each FOV. A colocalization test was then run between the two images from the same FOV to get a Pearson’s correlation value. For the imaging of the cells with 4 constructs expressed (one FLAG-tagged construct plus three constructs without FLAG tag), a 70-frame brief movie and a snapshot image were taken from the red (excitation: 561nm at 50 mW/mm^2^, emission: 579/34 nm) and far-red channel, respectively for each FOV. Four unmixed images were obtained via signal unmixing of each brief movie. The unmixed image from the FLAG-tagged construct was then co-localized with the image taken from the same FOV via far-red channel to get a Pearson’s correlation value. Cyan light (488 nm at 40 mW/mm^2^) was applied for 50 ms before brief movies were taken.

##### Crosstalk measurements

For the crosstalk measurements of green FPs, seeded NIH3T3 cells were transfected with pcDuex2-rsGreenF-E, pcDuex2-rsEGFP2-E, pcDuex2-rsFastLime, pcDuex2-Skylan62A, pcDuex2-Dronpa, and pcDuex2-YFP separately in 6 different wells of 24-well plates. Three hours after transfection, the growth medium with transfection reagents of each well was aspirated and the cells were then washed with 1x PBS three times followed by trypsin treatment (0.05% trypsin-EDTA (Gibco)) for 2 mins. The detached cells from each well were then suspended and collected before being fully mixed with the cells from the other 5 wells. Mixed cells were then seeded back to 24-well plates. 16-24 hours after re-seeding, imaging was performed on a Nikon Eclipse Ti inverted microscope equipped with a confocal spinning disk (CSU-W1), a 20×, 0.75 NA air objective, and a 5.5 Zyla camera (Andor), controlled by NIS-Elements AR software. For each FOV, a 70-frame brief movie was taken (exposure time, 50ms, excitation: 488nm at 10 mW/mm^2^, emission: 525/30 nm, on-switch:405 nm at 2.5 mW/mm^2^ for 100ms) and 6 unmixed images were obtained via signal unmixing. Since each transfected cell only expressed one FP, the crosstalks of the expressed FP to other FPs were calculated as the percentages of the fluorescence of other FP channels in that cell to the fluorescence of the expressed FP channel in the same cell.

The crosstalk measurements of red FPs were similar to that of green FPs except that only four plasmids (pcDuex2-rsTagRFP, pcDuex2-rsFusionRed1, pcDuex2-rScarlet, pcDuex2-mCherry) were used and brief movies were taken using the red channel (exposure time: 100ms, excitation: 561 nm at 12.5 mW/mm^2^, emission: 579/34 nm, on-switch: 488 nm at 10 mW/mm^2^ for 100ms).

#### TMI simulation

For simulations of temporal multiplexing, we used a pre-acquired fluorescent image (the image was taken on a Nikon epifluorescence inverted microscope with a 20×, 0.75 NA air objective; image size: 1024 × 1024) of NIH/3T3 cells expressing Dronpa to generate brief movies for both green FPs and red FPs. Segmentation was first applied to the fluorescent image to convert Dronpa-expressing cells to cell-shaped masks. Then, the normalized traces of six green FPs (or four red FPs) were scaled, each with a random ratio (the sum of the ratios equals 1) to create a hybrid trace for each mask (different masks contained different ratios of the six FPs, same masks contained the same ratios of the six FPs). Next, the values of hybrid traces (ranging from 0 to 1) at each time point (70 time-points in total) were used to multiply the fluorescence value of each pixel within the cell-shaped masks to generate a 70-frame brief movie (the fluorescence of the non-masking area was assigned as 0). Poisson noises were then calculated according to the fluorescence intensity at each pixel and then applied back to each pixel of the brief movie. In the meantime, six ground truth images (or four ground truth images for red FPs) were generated by multiplying the randomly assigned ratio of each FP by the fluorescence of the pre-acquired fluorescence image at each pixel within the cell-shaped masks. The fluorescence of the FPs in the non-masking area was assigned as 0. The code for TMI simulation is available on https://github.com/qiany09/Temporally-Multiplexed-Imaging.

#### Zebrafish Imaging

Procedures at MIT involving animals were in accordance with the National Institutes of Health Guide for the care and use of laboratory animals and approved by the Massachusetts Institute of Technology Animal Care and Use Committee. Zebrafish were raised and bred at 28 °C according to standard methods. DNA plasmids encoding rsTagRFP, rsFusionRed1, and rScarlet under the control of the 10x UAS promoter were mixed with a ratio of 2:2:1 and co-injected with Tol2 transposase mRNA into embryos of the pan-neuronal expressing Gal4 line, Tg(elavl3:GAL4-VP16) (*48*). Briefly, Tol2 transposase mRNA, synthesized using pCR2FA as a template (*49*) (mMESSAGE mMACHINE SP6 Transcription Kit, Thermo Fisher), and DNA was diluted to a final concentration of 25 ng/µl in 0.4 mM KCl solution containing 0.05% phenol red solution (Millipore Sigma) to monitor the injection quality. The mixture was kept on ice to minimize the degradation of mRNA during the injection. The mixture was injected into embryos at 1-cell stage (*50*). Larvae were screened for red fluorescence in the brain and spinal cord at day 3 post fertilization (animals were used without regard to sex) and subsequently imaged on day 5 post fertilization. To image zebrafish larvae, larvae were immobilized in 1.5% ultra-low-melting agarose (Millipore Sigma) prepared in E3 medium and paralyzed with 0.2 mg/ml pancuronium bromide (Millipore Sigma). Imaging was performed on a Nikon Eclipse Ti inverted microscope equipped with a confocal spinning disk (CSU-W1), a 40×, 1.15 NA water-immersion objective, and a 5.5 Zyla camera (Andor), controlled by NIS-Elements AR software. A 60-frame brief movie was taken for each FOV (exposure time 100ms, excitation: 561nm at 50 mW/mm^2^, emission: 579/34 nm, on-switch: 488 nm at 40 mW/mm^2^ for 50ms).

#### Imaging of cell cycle phases and kinase activities in NIH3T3 cells

FUCCI4 constructs were gifts from Michael Z Lin (Addgene no. 83841-83942) (*21*). Single-color FUCCI4 was built by replacing Clover, mKO2, mMaroon1, and mTurquoise2 with rsGreenFast-E, Dronpa, Skylan62A, and YFP respectively. 16-24 hours after transfection, NIH/3T3 cells were imaged on an epifluorescence inverted microscope (Eclipse Ti-E, Nikon) equipped with a 20×, 0.75 NA air objective, Perfect Focus System, an Orca-Flash4.0 V2 sCMOS camera (Hamamatsu), and a SPECTRA X light engine (Lumencor). Cells were placed in a stage-top incubator with a controlled environment at 37°C and 5% humidified CO_2_ (Live Cell Instrument), and brief movies (60 frames in 15 s, excitation light was on during the whole 15s) were acquired every 30 min. Excitation: 475/28 nm at 9.6 mW/mm^2^, emission: 525/50 nm, exposure time: 50ms, on switching: 390/22 nm at 1.2 mW/mm^2^ for 100ms.

The constructs of JNK KTR-rsGreenFast-E, P38 KTR-Dronpa, ERK KTR-rsFastLime were built based on the original KTRs constructs JNK KTR-Clover (Addgene plasmid 59151), P38 KTR-mCerulean3 (Addgene plasmid 59155), ERK KTR-Clover (Addgene plasmid 59150), all gift of Markus W. Covert (*22*). The newly developed three KTRs were then used along with PKA KTR-Clover (Addgene plasmid 59151) and H2B-TagBFP to report the activities of all four kinases. H2B-TagBFP was used as a nucleus marker. NIH/3T3 cells were imaged on the same microscope and incubator system as the imaging of cell cycle phases. Cell culture media were changed to imaging media (MEM (Gibco) without phenol red with 1% FBS (Gibco)) prior to imaging. The imaging conditions (including exposure time, excitation, and emission) for KTRs were the same as those for imaging cell cycle phases as described previously. Images of H2B-TagBFP (excitation: 390/22 nm at 1.2 mW/mm^2^; emission: 447/60 nm, exposure time: 100ms) were acquired right before recording brief movies to switch rsFPs to the “on”-state. Images and brief movies were acquired every 2 min. 50 μM Forskolin (Millipore Sigma), and 20 ng/ml basic fibroblast growth factor2 (bFGF2, R&D System) were used to activate kinase activities.

#### Combined temporal multiplexing and spectral multiplexing

##### Simultaneous imaging of cell cycle phases and activity of cyclin-dependant kinases

To increase the number of genes co-expressed within single cells, the four genes encoding FUCCI4 were cloned into a single plasmid as the following: CMV-rsGreenFast-Geminin_1-110_-P2A-Dronpa-cdt_30-120_-IRES-H1-Skylan62A-P2A-YFP-SLBP_18-126_. NIH3T3 cells were then transfected with the aforementioned plasmid, plasmid EF1α-DHB-TagBFP2 (*29*), and plasmid EF1α--mCherry-CDK4KTR (*30*). Imaging was performed 16-24 hours post-transfection on the same epifluorescence inverted microscope and incubation system as described previously. Blue channel (excitation: 390/22 nm at 1.2 mW/mm^2^; emission: 447/60 nm, exposure time: 100ms), green channel (60 frames in 15 s, excitation light (475/28 nm at 9.6 mW/mm^2^) was on during the whole 15s, emission: 525/50 nm; exposure time: 50ms) and red channel (excitation: 555/28 at 9.4 mW/mm^2^, emission: 630/75 nm; exposure time: 100ms) were used together for imaging of 6 signals. Purple light illumination used in the blue channel for excitation also served as the “on” trigger for green rsFPs. Images and brief movies were acquired every 30 min for 24-48 hours without stimulation.

##### Simultaneous imaging of seven cell activities within single cells

The genes of the single-color KTRs were cloned into the following plasmid: CAG-JNKKTR-rsGreenF-E-P2A-ERK KTR-rsFastLime-IRES-PKAKTR-Clover-P2A-P38KTR-Dropna. The aforementioned plasmid was then used with plasmid CAG-NIR-GECO2G (*33*), plasmid CMV-Pink Flamindo (*34*), and plasmid CMV-BlueCKAR (*2*) for imaging 7 signals within individual NIH/3T3 cells. Imaging conditions for blue channel, green channel, and red channel were the same as those used in the imaging of cell cycle phases and CDK activities. An extra channel (excitation: 637/12 nm at 9 mW/mm^2^; emission: 664LP, exposure time: 100ms) was used for imaging NIR-GECO2G. Images and brief movies were acquired every 2 min. 50 μM forskolin (Millipore Sigma) and 100 ng/ml phorbol 12-myristate 13-acetate (PMA) (Millipore Sigma) were used to stimulate cells.

#### Signal unmixing and image analysis

##### Signal unmixing of temporal multiplexed imaging

Reference traces of FPs used for signal unmixing were collected right before or after each imaging experiment. Each reference trace was an averaged result from 10 to 30 cells of 2 to 3 brief movies from one cell culture batch. Traces from each cell were normalized to the maximum value before averaging. Cells were selected so they were evenly distributed on the fields-of-views. For signal unmixing of temporal multiplexing imaging, the recorded trace at each pixel was first normalized to the maximum value and then unmixed into a linear combination of the reference traces of fluorophores using least squares regression. Next, the resultant ratios (ranging from 0 to 1) of fluorophores at each pixel were multiplied by the fluorescence value of the first frame of the brief movie at this very pixel to generate unmixed images. The code for signal unmixing is available on https://github.com/qiany09/Temporally-Multiplexed-Imaging.

Brief movies were processed in Fiji as follows before being subjected to signal unmixing using custom Matlab code (or before being used for extracting reference traces): images were down-sampled from size 2048×2048 to size 1024×1024 or size 512×512 (to decrease computing time) followed by background subtraction.

##### Quantification of KTRs

Kinase activities reported by KTRs including the CDK2 reporter and CDK4/6 reporter were quantified following the methods described previously (*22*). Briefly, to calculate cytoplasmic intensity to nuclear intensity, a nucleus and a five-pixel-wide cytoplasm ring were segmented for each cell via nucleus-targeted fluorescent proteins. Nucleus segmentation and cell tracking were performed in Fiji using StarDist (*51*) and trackmate, respectively. Cytoplasmic rings were segmented by using a custom macro in Fiji. Median intensity extracted from each ROI was used to calculate ratios. The ratios reflect kinase activity. CDK4/6 activity was then corrected by deducting 0.35-fold CDK2 activity, as before (*30*).

##### Analysis of CDK2 and CDK4/6 activity traces (Fig. 5 C, D)

CDK traces of each cell were first divided into sub-traces of different cell cycle phases. Each sub-trace of a complete cell cycle phase was then evenly split into three parts. A value was obtained by averaging all the data points in one part, thus yielding 3 values representing CDK activity at early, middle and late stages of each cell cycle phase, for a given cell.

All images in the manuscript were processed and analyzed using Fiji. Traces and graphs were generated using GraphPad prism 8 or Origin9.0.

## Supplementary Figures

**Fig. S1.**
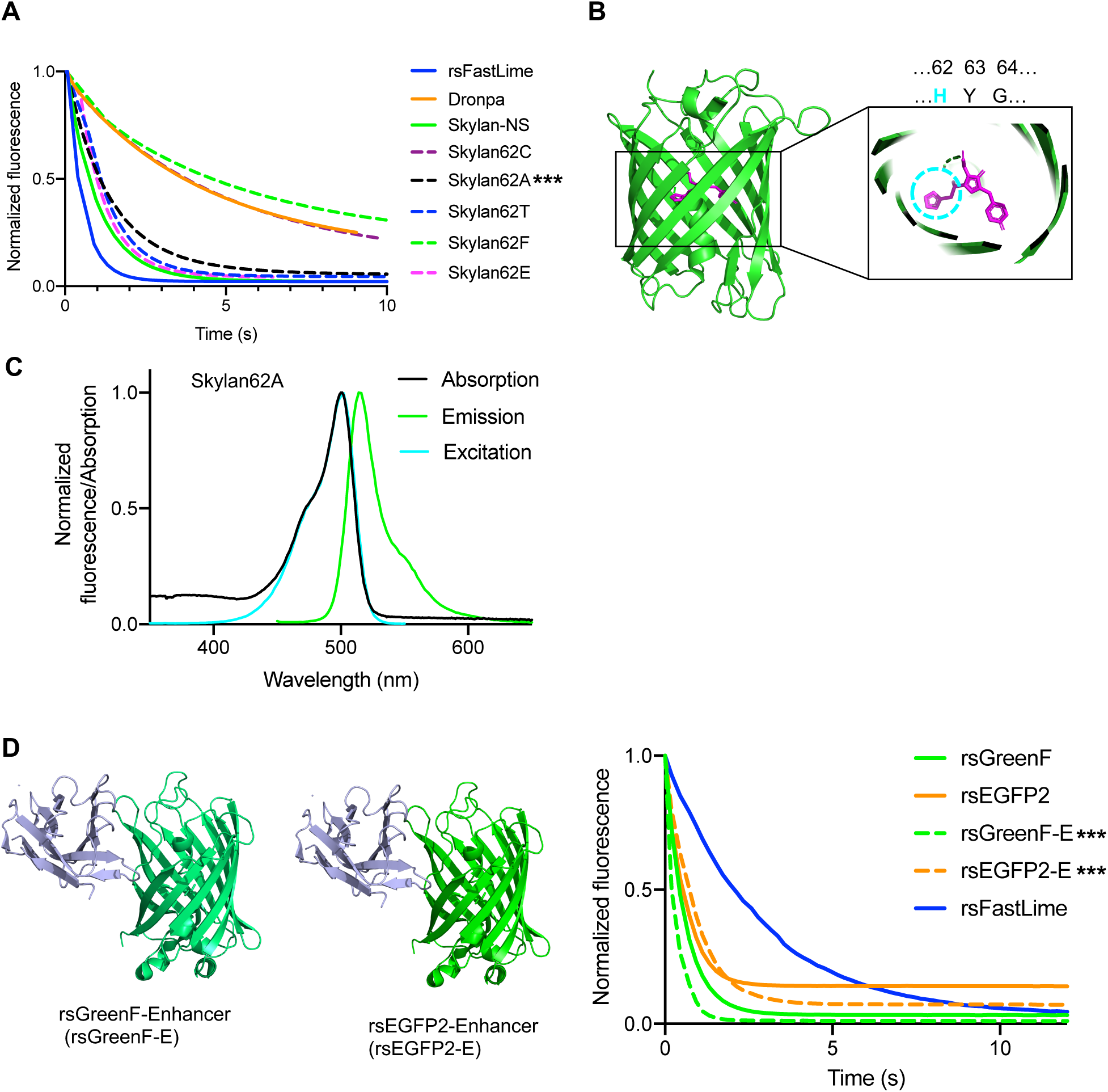
Development of new green rsFPs for temporally multiplexed imaging. (**A**) Off-switching traces of a series of Skylan mutants obtained in HEK293FT cells; rsFastLime and Dronpa were also tested under the same conditions for reference. Averaged values are shown, n = 8 - 12 cells from one culture. Excitation: 475/28 nm at 15 mW/mm^2^. Skylan62A (indicated by triple stars) was the winner of the screening. (**B**) Crystal structure of mEos (PDB: 3S05). The chromophore (highlighted in magenta) of mEos is formed by the tripeptide HYG. Inspired by the engineering of photoswitchable Skylan-NS from mEos3.1, which was achieved by mutating His62 (highlighted by dashed cyan circle) to Leu, a new photoswitchable FP, Skylan62A was developed from Skylan-NS by mutating Leu 62 to Ala. The off-switching rate of Skylan62A is in between that of Dronpa and rsFastLime, which makes Skylan62A a good candidate for temporally multiplexed imaging with rsFastLime and Dronpa. (**C**) Absorption (black), excitation (cyan), and emission (green) spectra of Skylan62A. (**D**) Left, structure of rsGreenF-Enhancer (rsGreenF-E) and rsEGFP2-Enhancer (rsEGFP2-E). Right, off-switching traces of rsGreenF and rsEGFP2 with and without enhancer recorded in HEK293FT cells. rsFastLime was also tested under the same conditions for reference. Averaged values are shown, n = 6 - 10 cells from one culture. Excitation: 475/28 nm at 5 mW/mm^2^. The addition of a nanobody enhancer increases the photoswitching kinetics of rsGreenF and decreases the photoswitching rate of rsEGFP2. rsGreenF-E and rsEGFP2-E (indicated by triple stars) were then chosen for temporally multiplexed imaging.

**Fig. S2.**
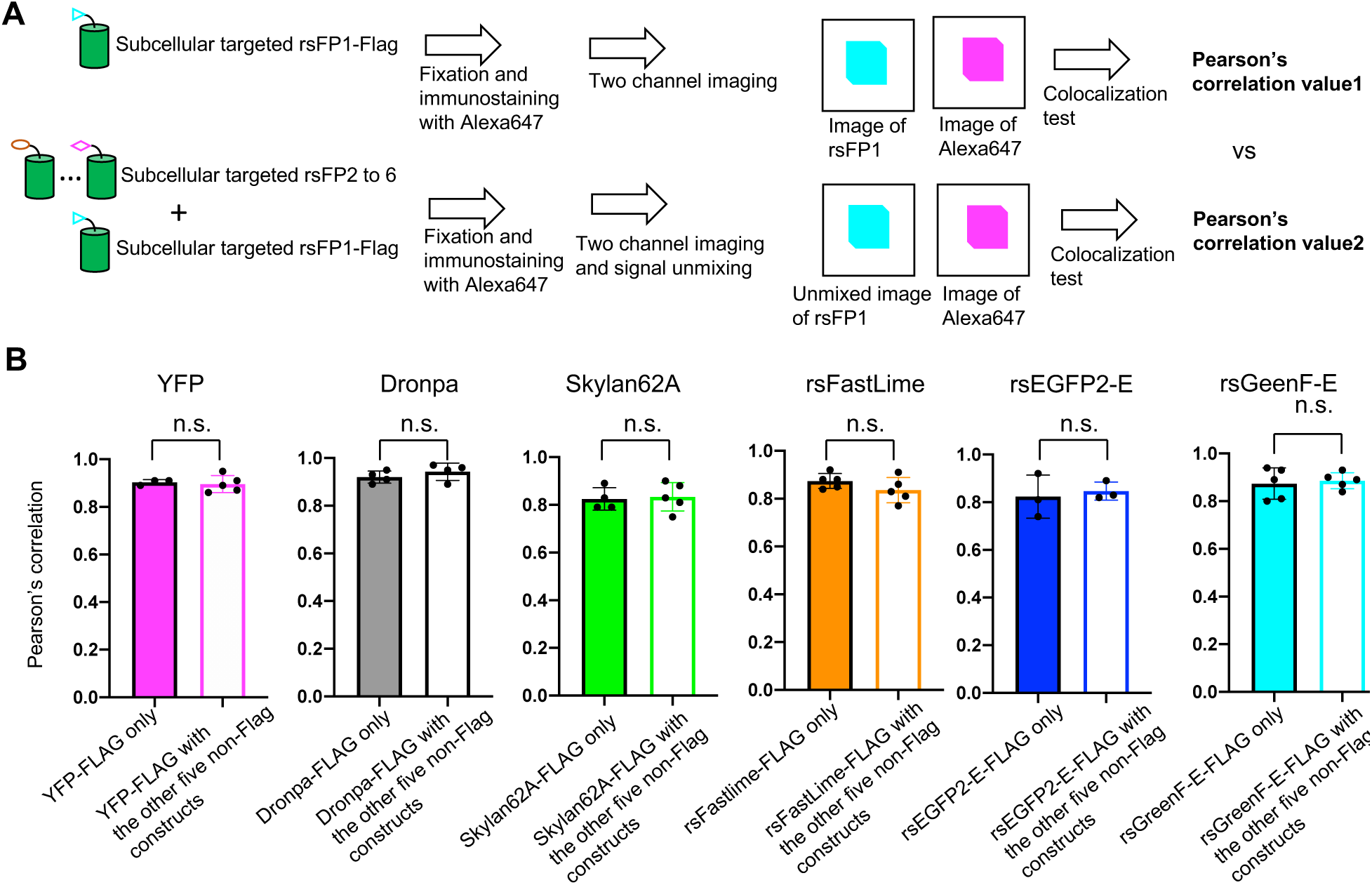
Evaluation of TMI for subcellular labeling with six green FPs. (**A**) Experimental design for evaluating temporal multiplexed imaging using green FPs. A FLAG tag was added to each of the six subcellularly targeted constructs. In the first experiment, as shown on the upper panel, each FLAG-tagged construct was expressed individually in U2OS cells. 48 hours post transfection, cells were fixed and immuno-stained with Alexa647 dye. Two snapshot images were taken from green channel (excitation: 488nm, emission: 525/30 nm) and far-red channel (excitation: 637nm, emission: 700/50nm) followed by colocalization test and calculation of Pearson’s correlation values. In the second experiment, as shown on the bottom panel, one FLAG-tagged construct was coexpressed with other five constructs without a FLAG tag in U2OS cells, 48 hours post transfection; a 70-frame brief movie was taken from the green channel and a snapshot image was taken from the far-red channel. After signal unmixing, six images were obtained from the green channel. The unmixed image of the FLAG-tagged construct was then co-localized with the image obtained from far-red channel followed by calculation of another set of Pearson’s correlation values. The two sets of Pearson’s correlation values were then compared for each FP. (**B**) Bar plots of Pearson’s correlation values of each FP obtained in **A**. n = 3 to 6 images from two cell culture batches. Data are shown as mean ± SD with individual values plotted as dots. Unpaired t test was run between two sets of Pearson’s correlation values for each FP; n.s., no significance.

**Fig. S3.**
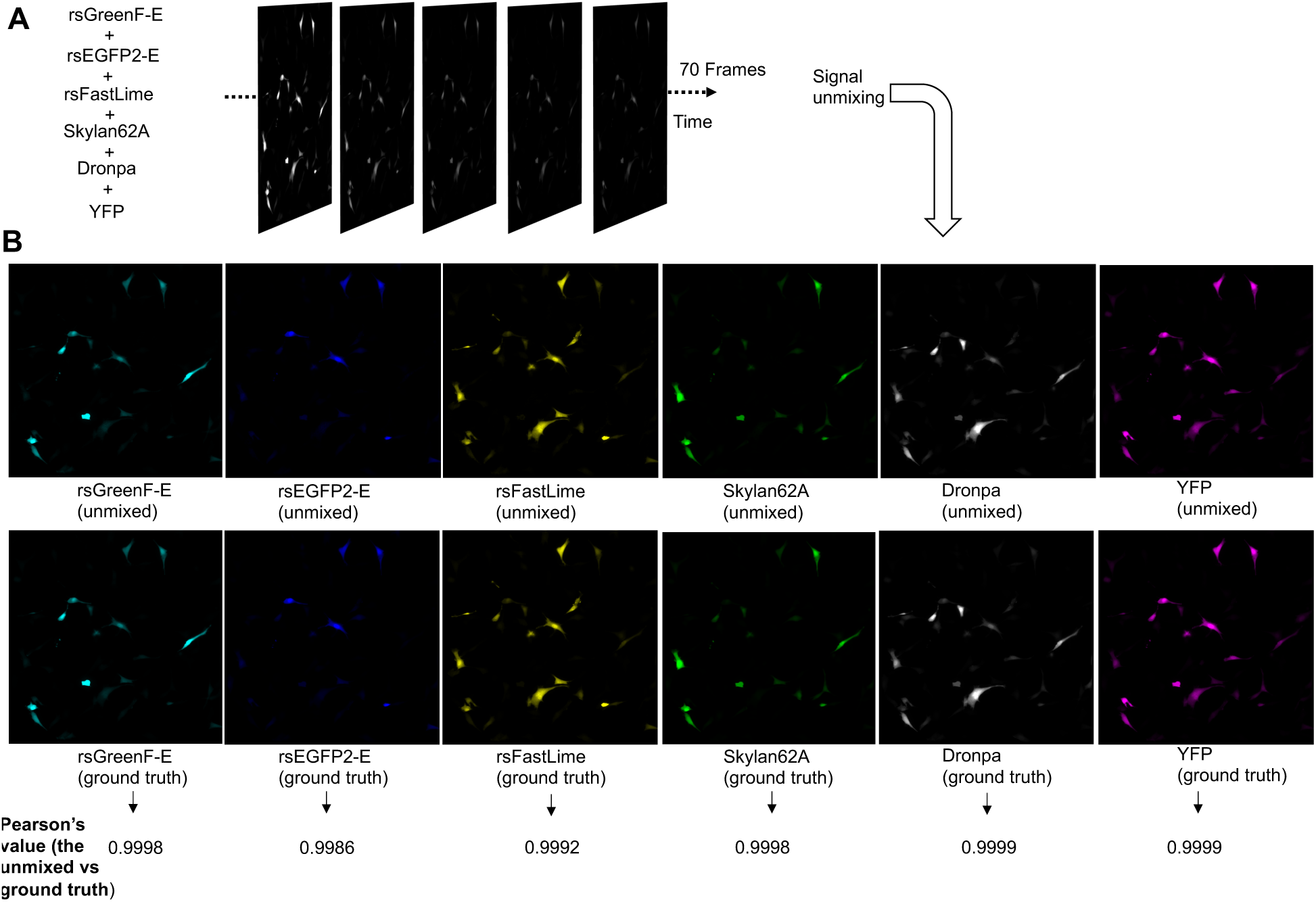
Simulation of temporally multiplexed imaging of six green FPs. A 70-frame brief movie was generated using a pre-acquired fluorescence image of NIH/3T3 cells and the off-switching traces in **Fig. 2B**. The fluorescence components of the six FPs in each cell were randomly assigned. Different cells, thus, had different fluorescence combinations of the six FPs. Poisson noise was added to each pixel to mimic real images. (**A**) Simulated brief movie showing cells that express six green rsFPs. (**B**) After signal unmixing, six images were obtained for the FPs. The images were then compared with ground-truth images. Pearson’s correlation values between unmixed images and ground-truth images are shown. Detailed value comparisons are shown in **Fig. 2F**.

**Fig. S4.**
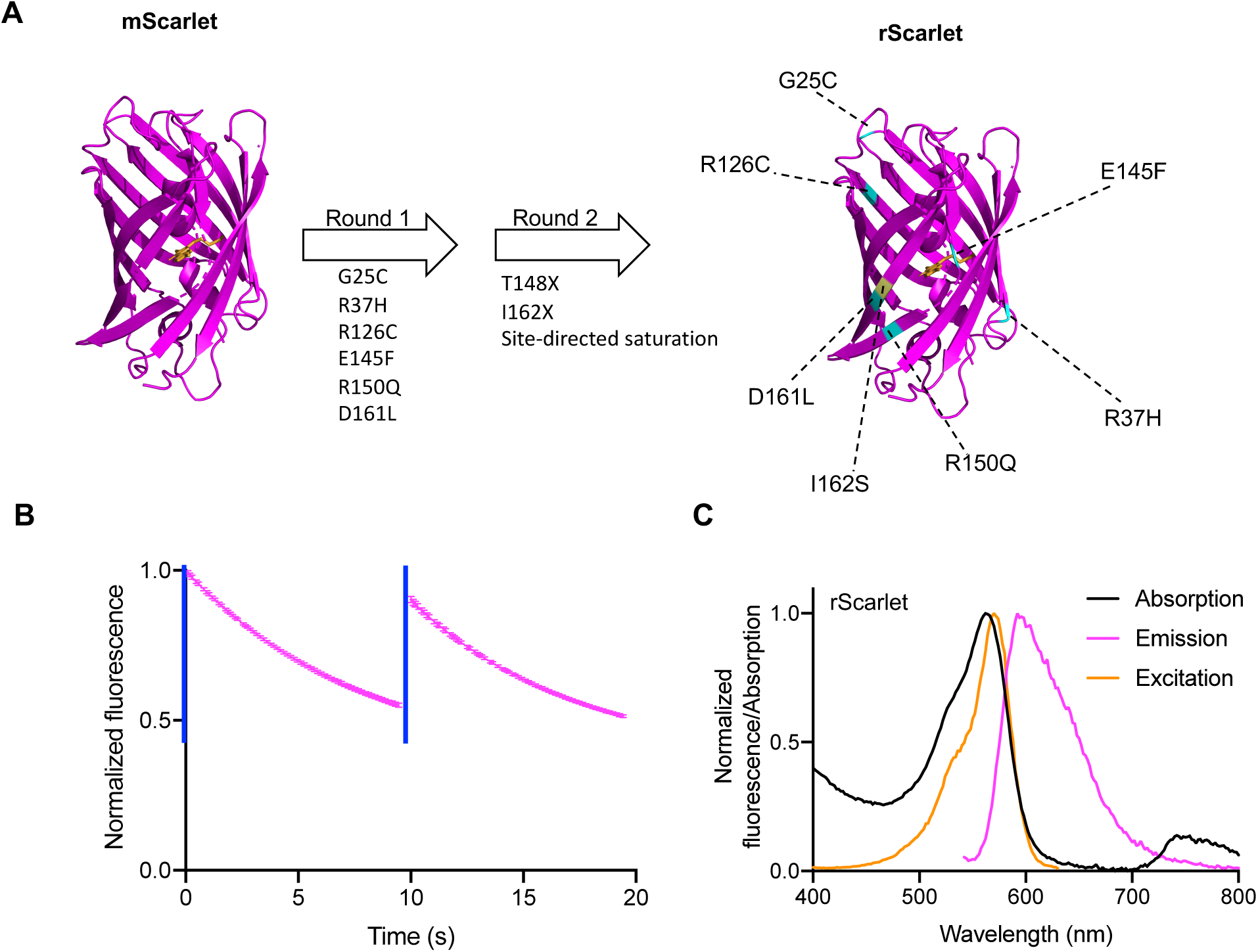
Development of a new red rsFP with a slow photo-switching rate for temporally multiplexed imaging. (**A**) Schematic illustration of the engineering of rScarlet from mScarlet. Left, crystal structure of mScarlet; the chromophore is shown in orange. Two rounds of evolution were performed before photoswitchable rScarlet was selected. rScarlet accumulated 7 mutations compared to mScarlet, which is highlighted in the crystal structure on the right. The mutations from round 1 are highlighted in cyan, and the mutation (I162S) from round 2 is highlighted in yellow (the amino acid at site no. 148 didn’t change during the screening). (**B**) Photoswitching traces of rScarlet in HEK293FT cells, n = 3 cells from one culture. Data are shown as mean ± SD. Excitation: 555/28 nm at 9.4 mW/mm^2^, on-switching: 475/28nm at 9.6 mW/mm^2^ for 100 ms (blue bar). (**C**) Absorption (black), excitation (orange), and emission (magenta) spectra of rScarlet.

**Fig. S5.**
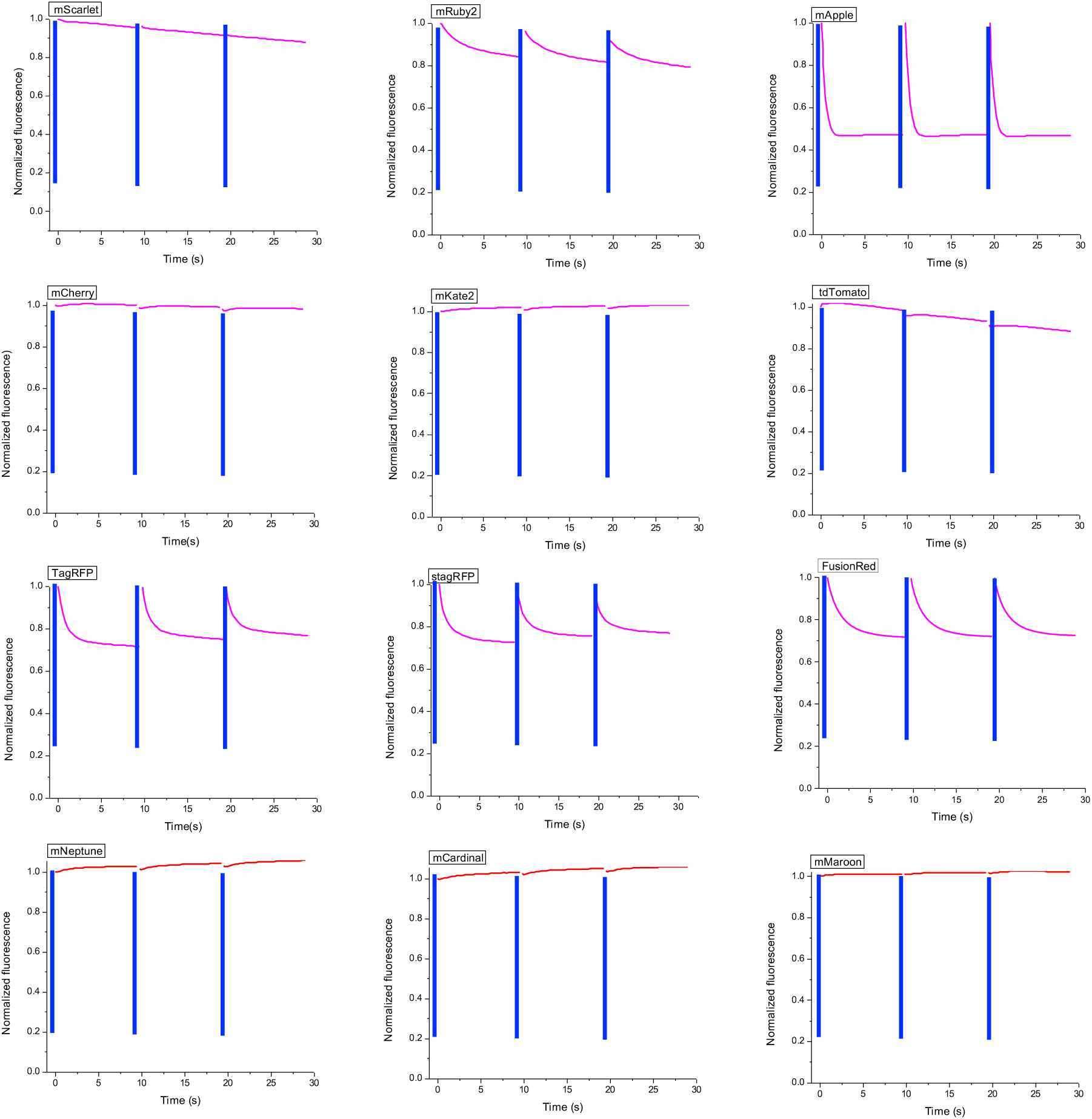
Photoswitching tests of commonly used red (magenta) and far-red (red) fluorescent proteins in HEK293FT cells. Averaged values are shown, n = 6 - 12 cells from one cell culture batch. Excitation: 555/28 nm at 9.4 mW/mm^2^; on-switching: 475/28 nm at 9.6 mW/mm^2^ for 50 ms (blue bar).

**Fig. S6.**
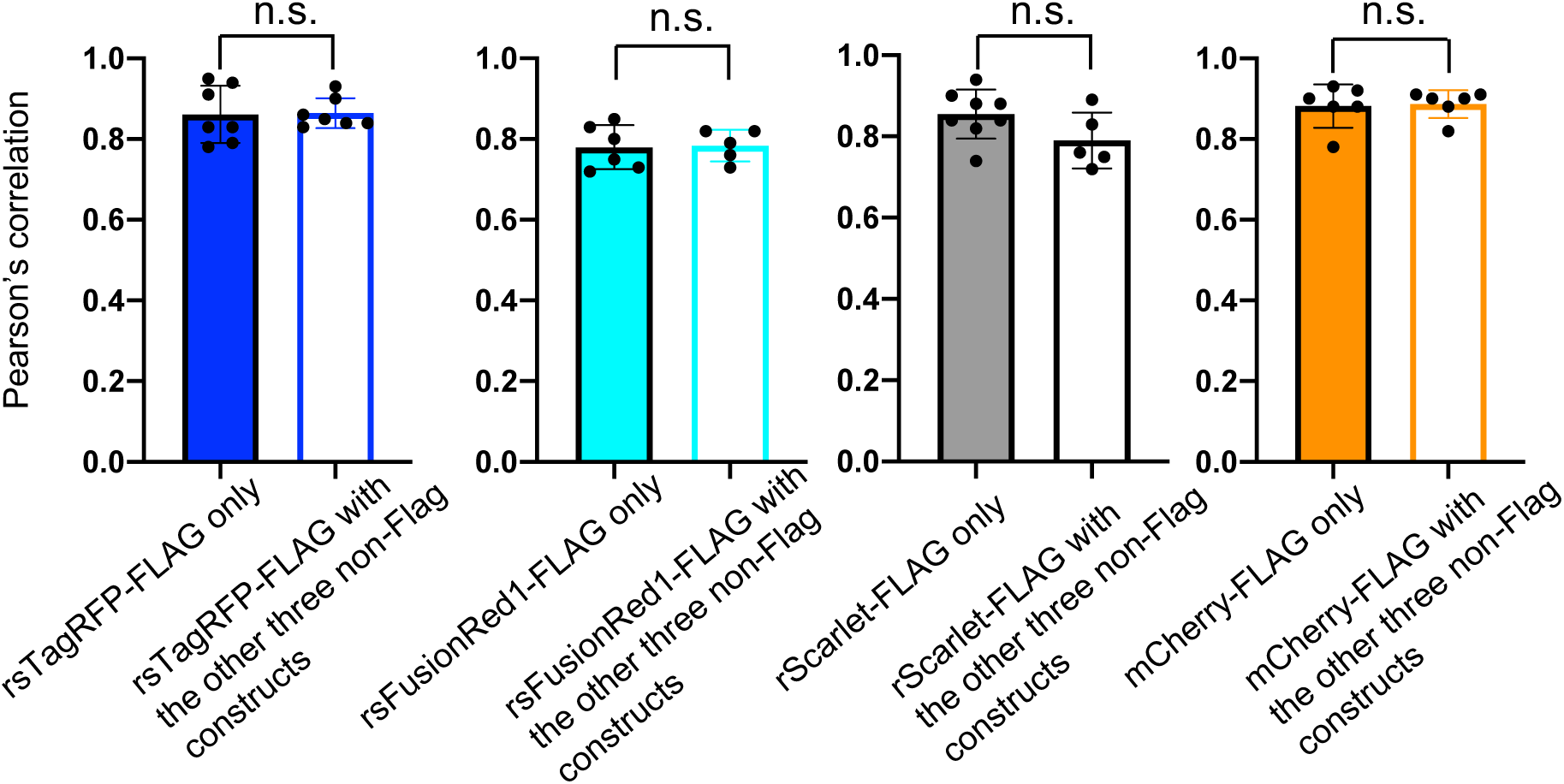
Evaluation of TMI for subcellular labeling with four red FPs. The same experiments as in **Fig. S2** were run for the four red FPs used in TMI, and two sets of Pearson’s correlation values were obtained for each red FP. Bar plots of mean with SD are used, with individual values plotted as dots; n = 5 to 6 images from two cell culture batches. Unpaired t-tests were run between two sets of Pearson’s correlation values for each FP; n.s., no significance.

**Fig. S7.**
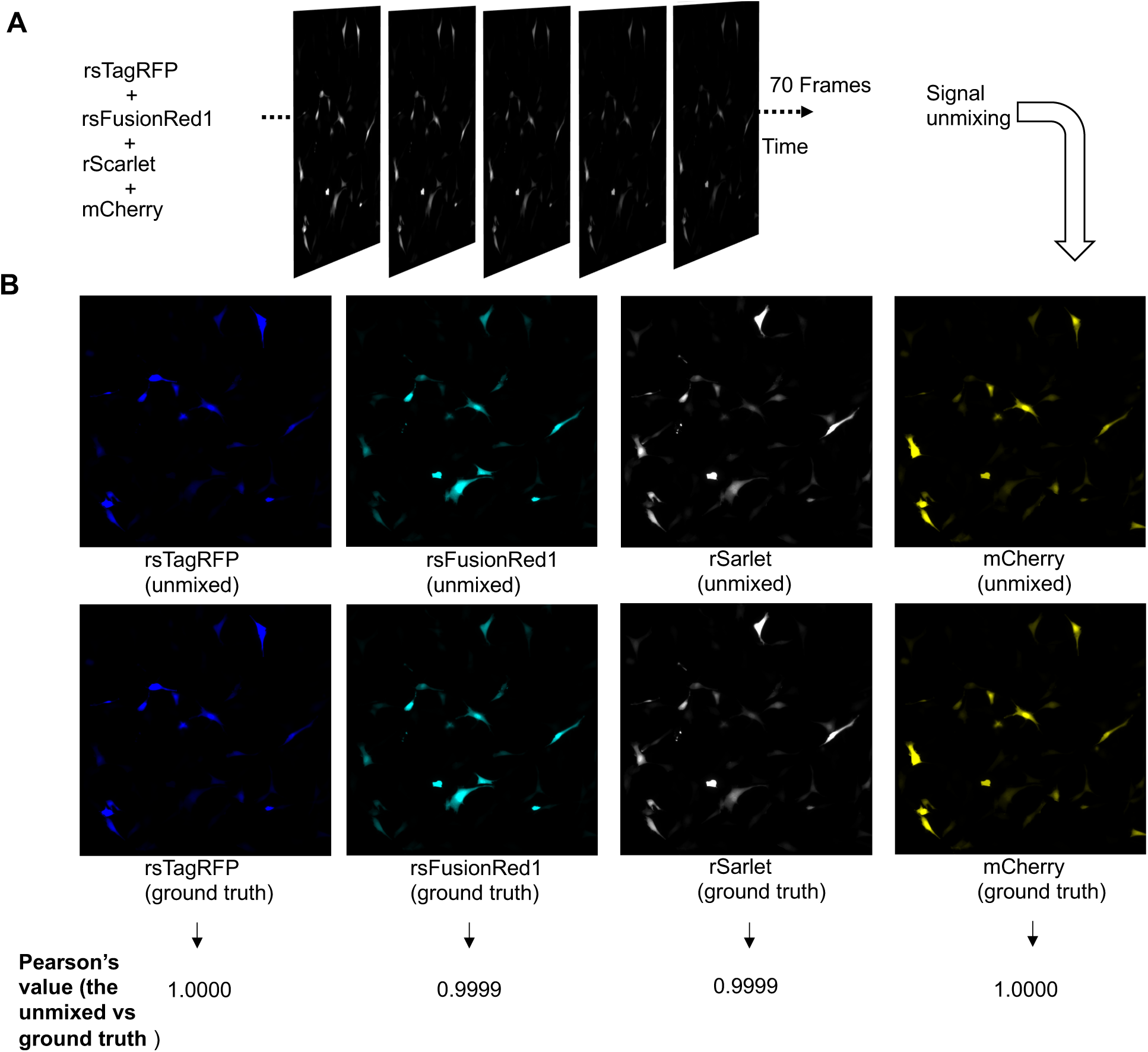
Simulation of temporally multiplexed imaging of four red FPs. A 70-frame brief movie was generated using a pre-acquired fluorescence image of NIH/3T3 cells (same image as used in **Fig. S3**) and the off-switching traces in **Fig. 3B**. The fluorescence components of the four red FPs were randomly assigned and mixed. Different cells had different fluorescence combinations of the four red FPs. Poisson noise was added to each pixel to mimic real images. (**A**) Simulated brief movie showing cells that express four red FPs. (**B**) After signal unmixing, four images were obtained for the FPs. The images were then compared with ground-truth images. Pearson’s correlation values between unmixed images and ground-truth images are shown. Detailed value comparisons are shown in **Fig. 3F**.

**Fig. S8.**
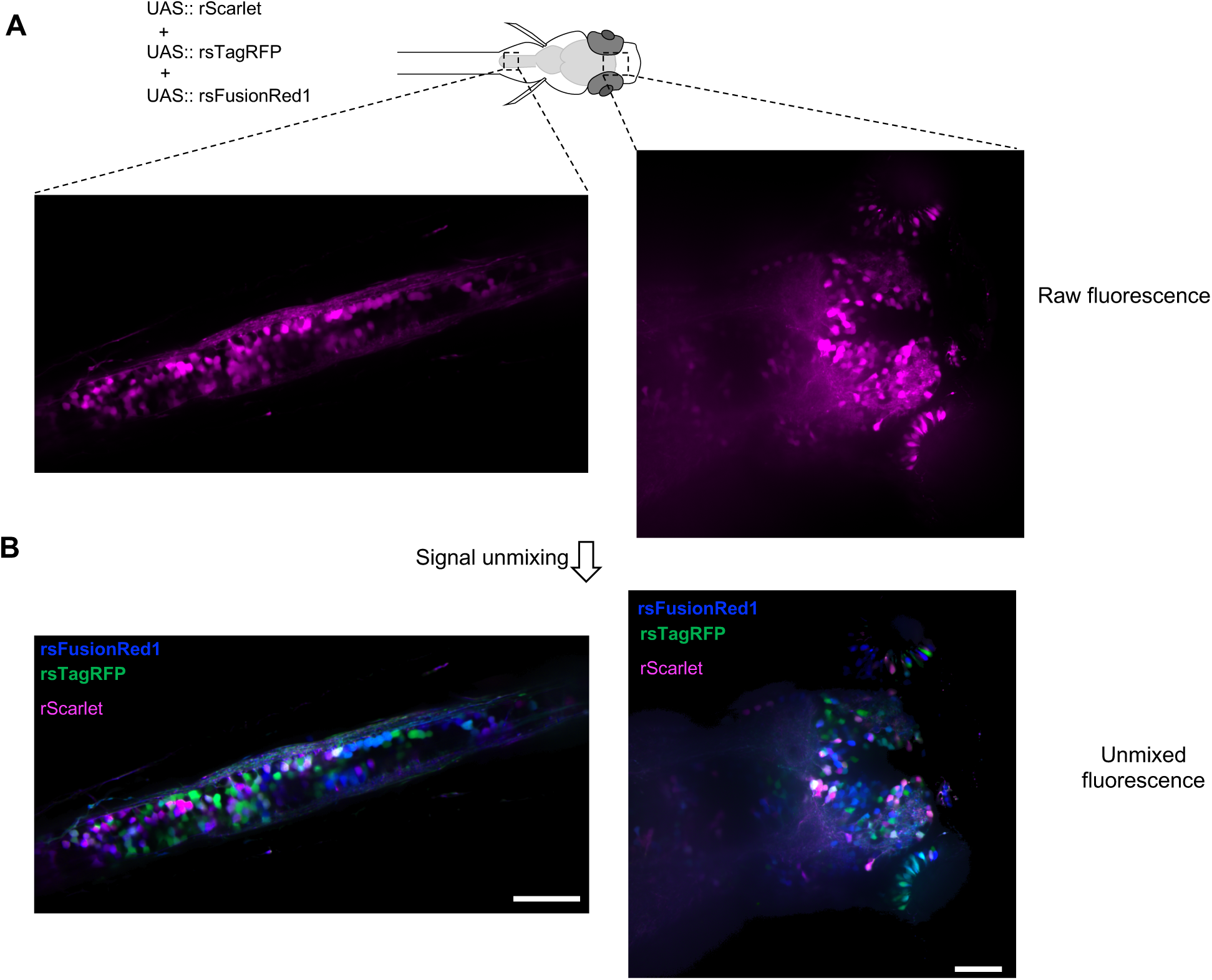
More images of single-color “brainbow” in zebrafish. (**A**) Representative raw fluorescent images (out of 20 images taken from 3 animals) of zebrafish larvae in spinal cord and forebrain (dorsal view). (**B**) Brainbow-like images showing the same area in **A**. Scale bars, 20 µm.

**Fig. S9.**
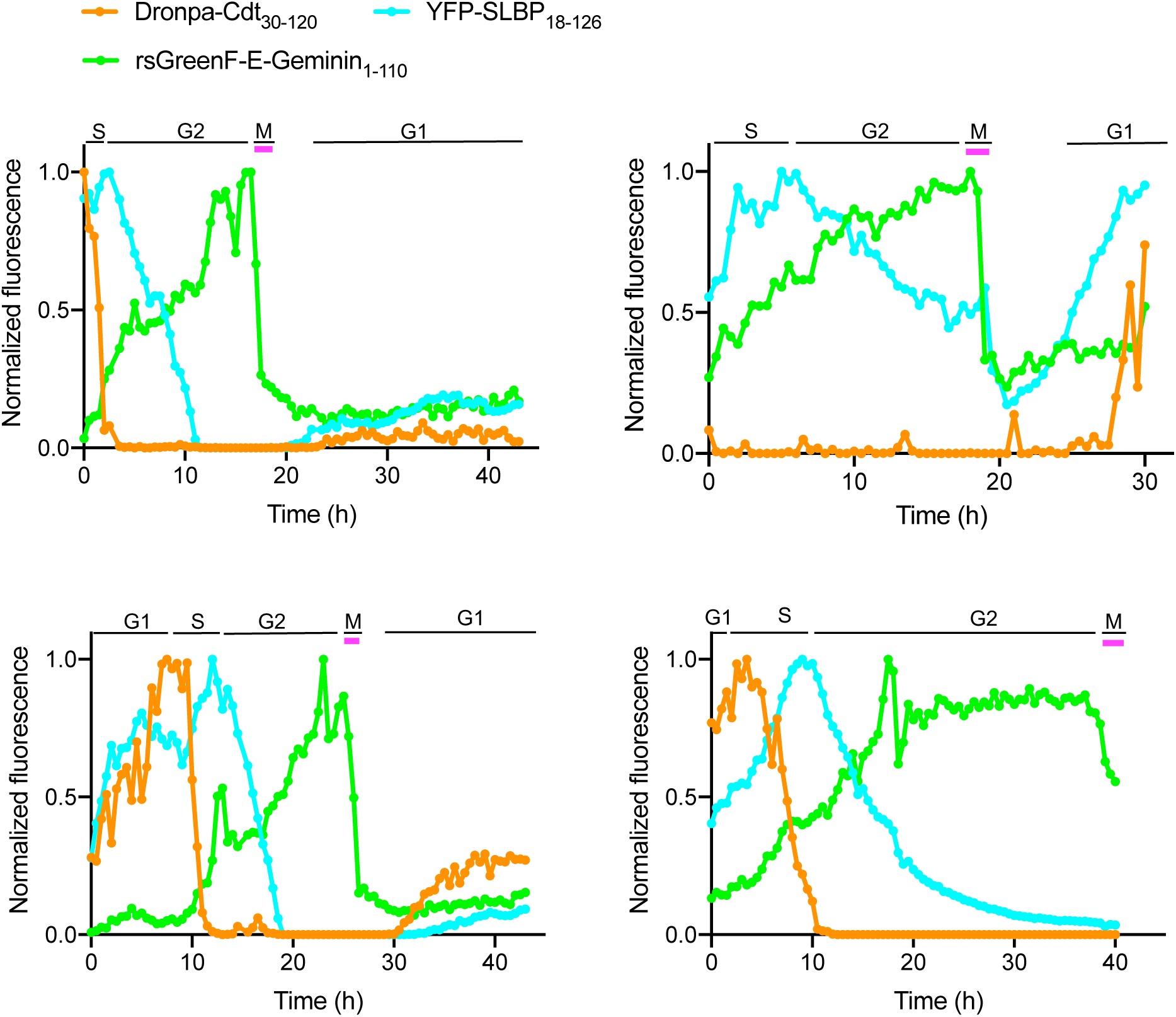
Additional fluorescence traces of Dronpa-Cdt1_30-120_ (orange), rsGreenF-E-Geminin_1-110_ (green), and YFP-SLBP_18-126_ (Cyan) during cell divisions, tracing one arbitrary daughter cell from each mother cell when the cell divides. Fluorescence was normalized to maximum value. Magenta bars indicate observation of chromosome condensation. Cell-cycle phases were assigned based on the principles in **Fig. 4A**.

**Fig. S10.**
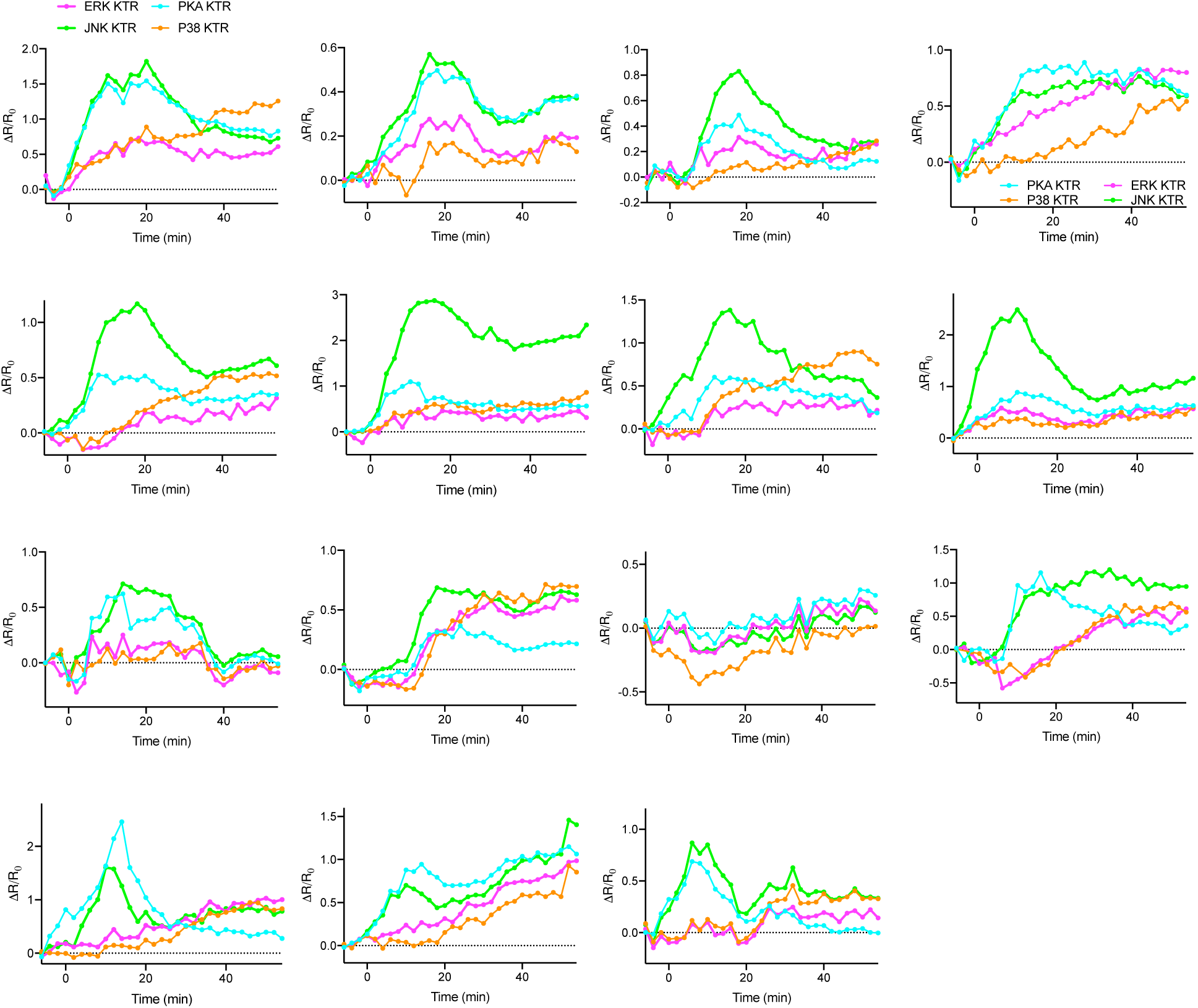
Activity traces of ERK KTR-rsFastLime, JNK KTR-rsGreenF-E, P38 KTR-Dronpa, PKA KTR-Clover from individual cells upon stimulation with 20 ng/ml bFGF2, added at t = 0 min. The averaged traces are shown in **Fig. 4F**. The same color code is used throughout the figure. R: cytoplasmic intensity to nuclear intensity ratio. Change in fluorescence of the sensors is plotted as ΔR/R_0_ (R_0_ was the averaged value from t = −6 min to t= −4 min).

**Fig. S11.**
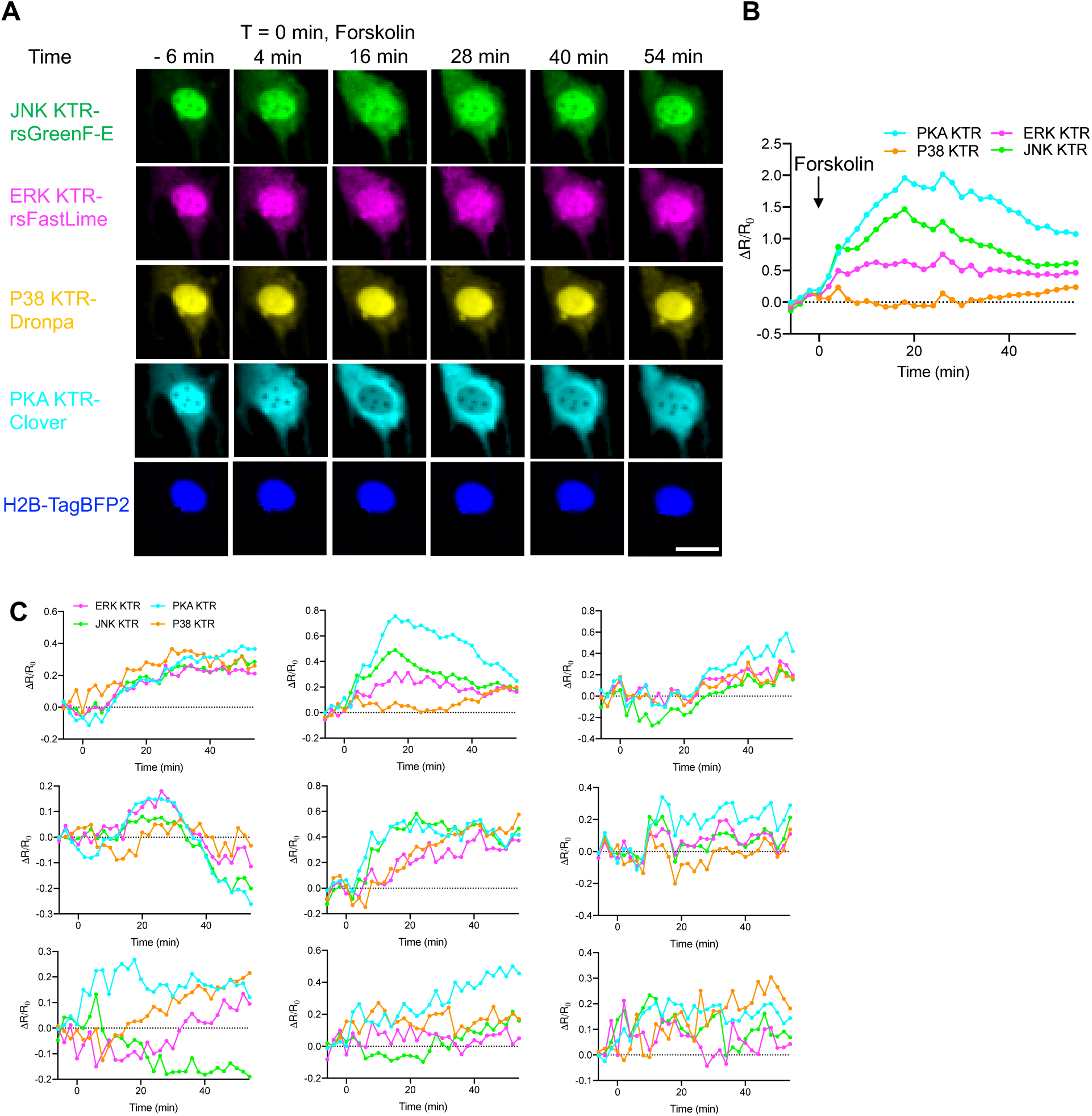
Responses of ERK KTR-rsFastLime, JNK KTR-rsGreenF-E, P38 KTR-Dronpa, PKA KTR-Clover after treatment with forskolin. (**A**) Represented NIH/3T3 cell expressing all four KTR sensors and H2B-TagBFP was stimulated with 50 µM forskolin and imaged at the indicated time points. Forskolin was added at t = 0 min. Scale bar, 20 µm. (**B**) Fluorescence traces of ERK KTR, JNK KTR, P38 KTR, and PKA KTR from the cell in **A**. R: cytoplasmic intensity to nuclear intensity ratio. Change in fluorescence of the sensors is plotted as ΔR/R_0_ (R_0_ was the averaged value from t = −6 min to t= −4 min). (**C**) Additional kinase activity traces from individual cells. Forskolin was added at t = 0 min. The same color code is used for all the traces throughout the figure.

**Fig. S12.**
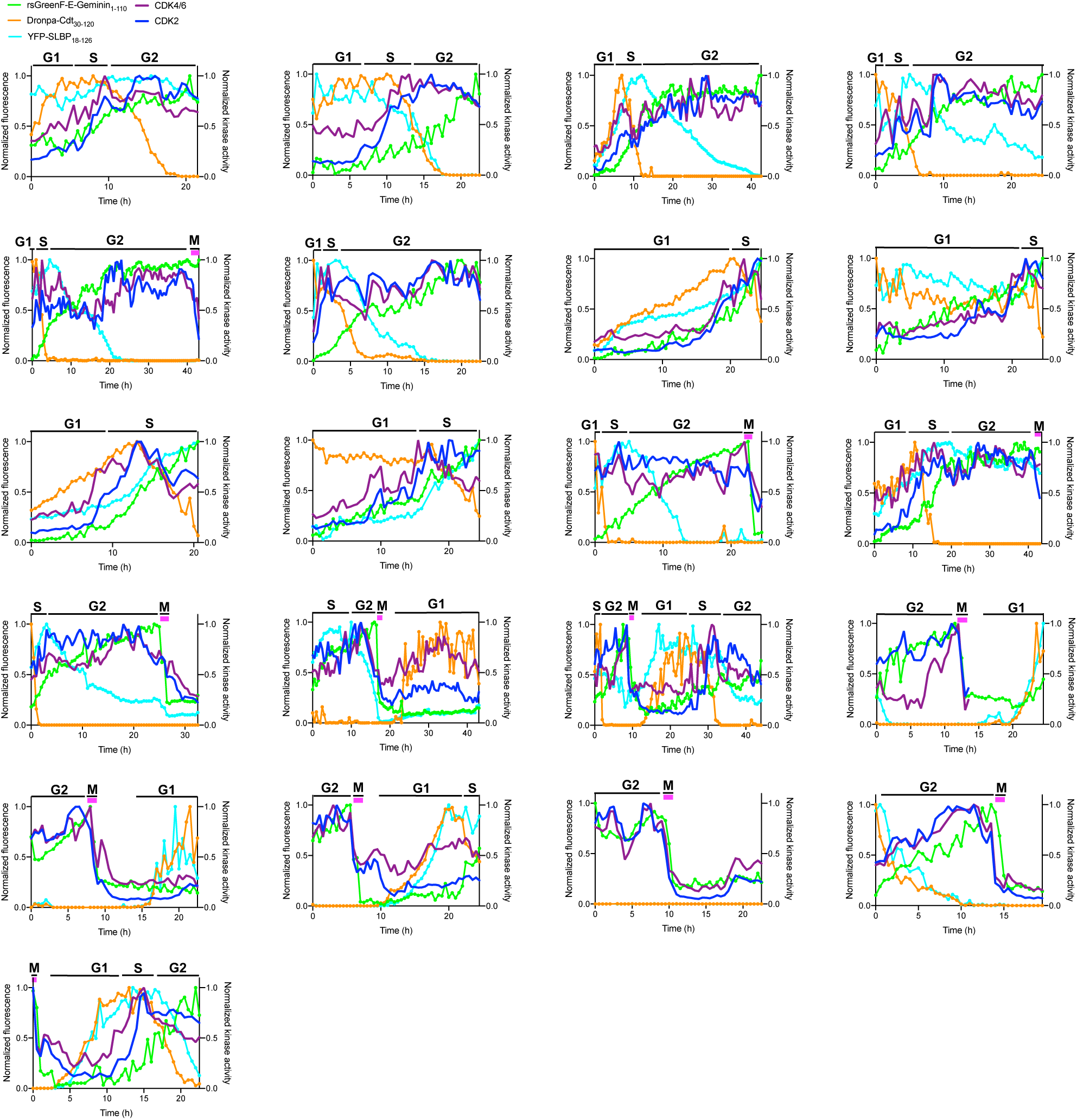
Additional traces of four fluorescence signals from FUCCI4 and two CDK signals from corresponding reporters over cell divisions. NIH/3T3 cells were imaged every 30 min without stimulation. Fluorescence signals of Dronpa-Cdt1_30-120_ (orange), rsGreenF-E-Geminin_1-110_ (green), YFP-SLBP_18-126_ (Cyan), CDK2 reporter (blue) and CDK4/6 reporter (magenta) were calculated and shown in the same ways as described in **Fig. 5B**.

**Fig. S13.**
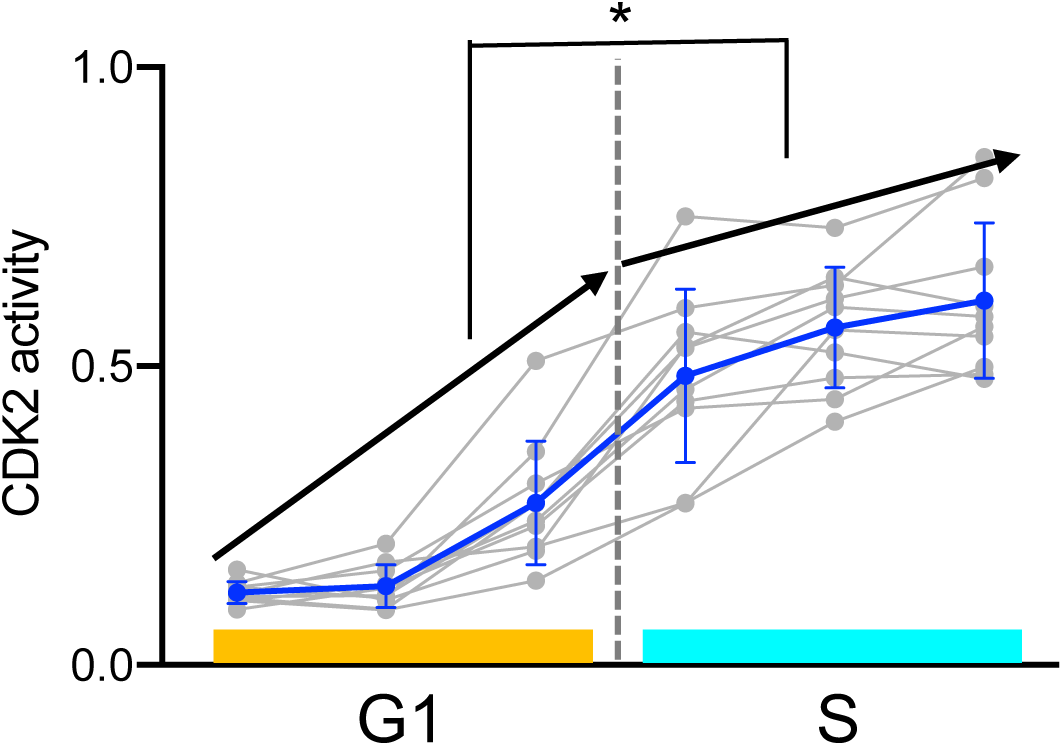
Comparison of CDK2 activity accumulation between G1 and S phase. Data are shown as mean ± SD (blue) with all individual values plotted (grey). Wilcoxon rank sum test was run between the difference in CDK2 activity between the end and beginning of G1 and the difference in CDK2 activity between the end and beginning of S phase, n = 10 cells from 5 cell culture batches. Wilcoxon rank sum tests, *p < 0.05.

**Fig. S14.**
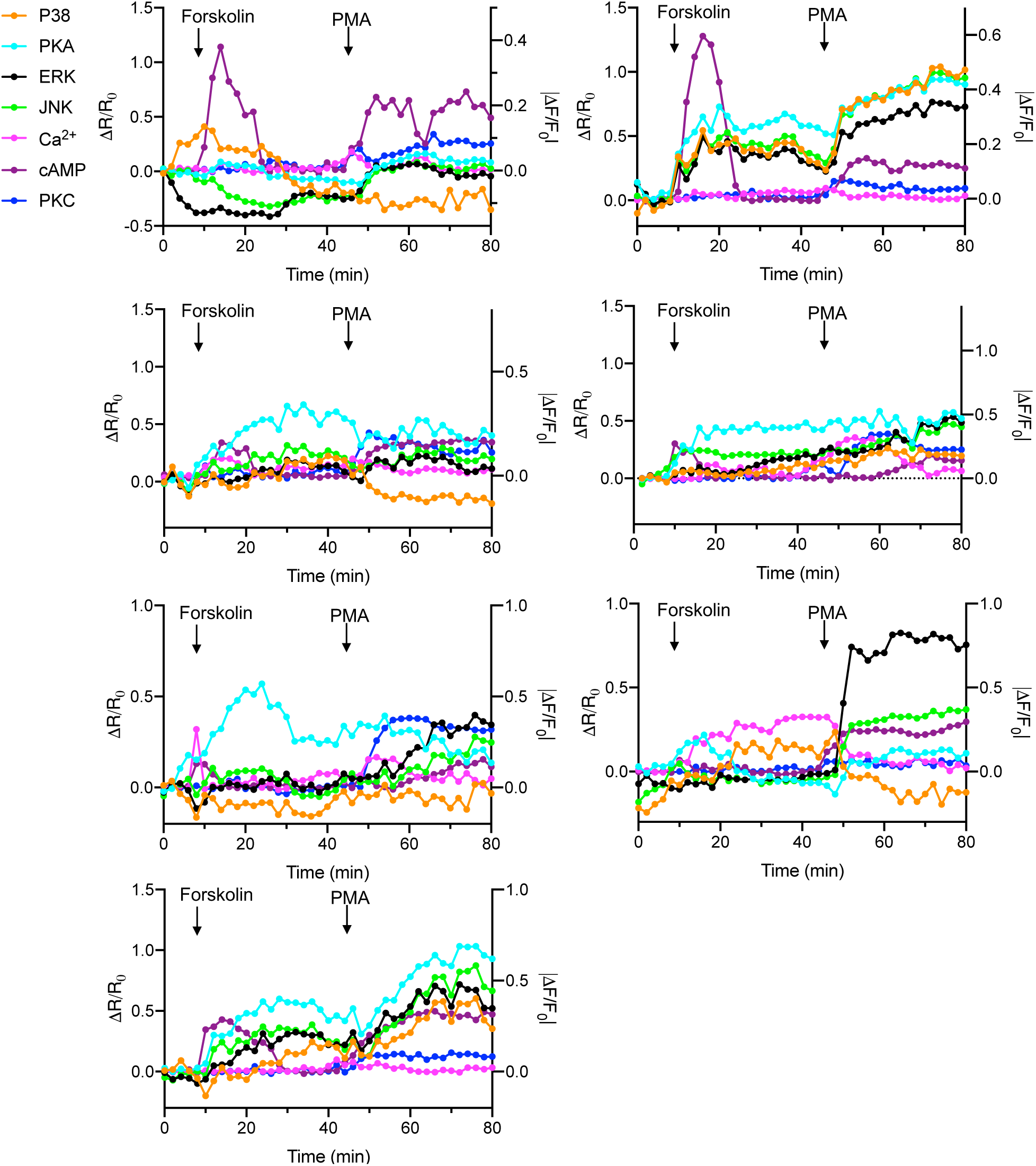
Imaging of 7 signals from individual NIH/3T3 cells. Activity traces of P38 (orange), PKA (cyan), ERK (black), JNK (green), Ca^2+^ (magenta), cAMP (purple), and PKC (blue) from single cells.

**Fig. S15.**
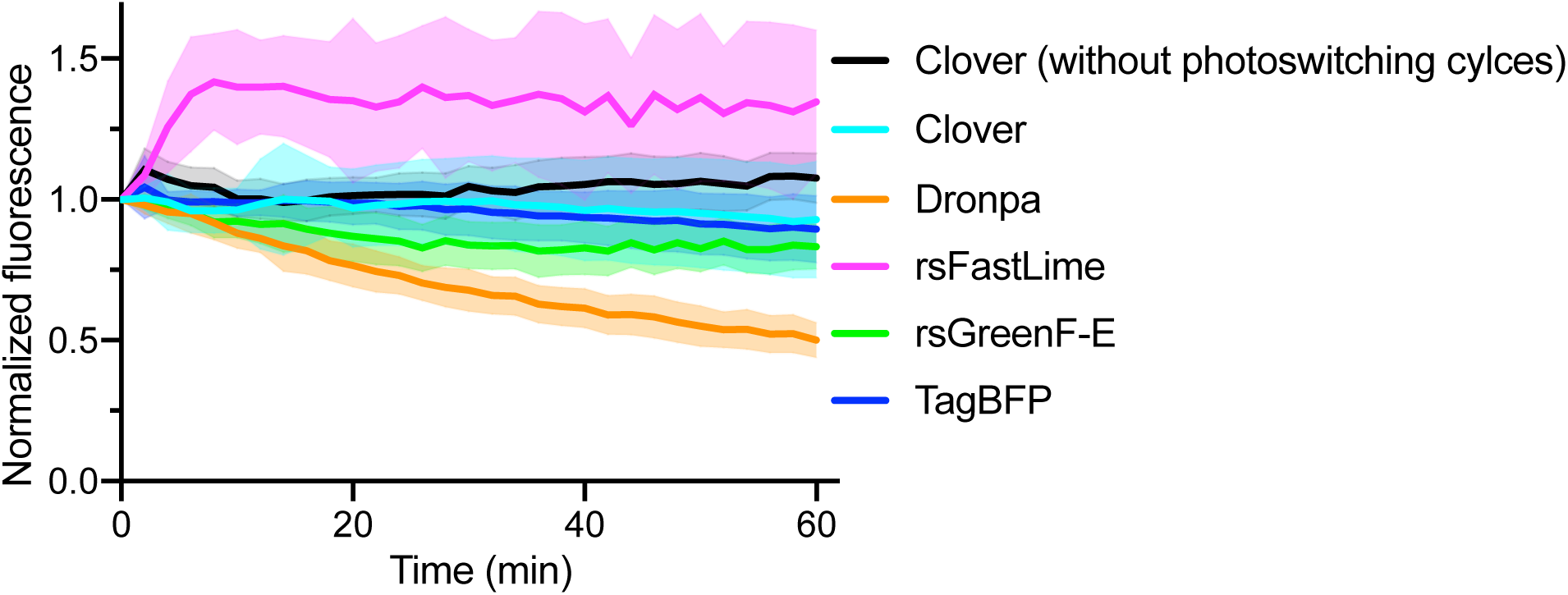
Photobleaching profiles of green FPs used in TMI for imaging of kinase activity. Data are extracted from the unmixed images obtained in the experiments in Fig. 4E-H, Fig. S10 and Fig. S11 (images of TagBFP are directly from the blue channel). Data are shown as mean ± SD, n = 23 cells from 4 cell culture batches. Clover was also tested in a separate experiment without taking brief movies (only one snapshot image was taken every 2 min; exposure time, 50ms; imaging conditions were the same as those in Fig. 4E-H, Fig. S10 and Fig. S11), serving as a reference for conventional imaging; n = 7 cells from 1 cell culture batch.

## Supplementary Tables

**Table Sl.**
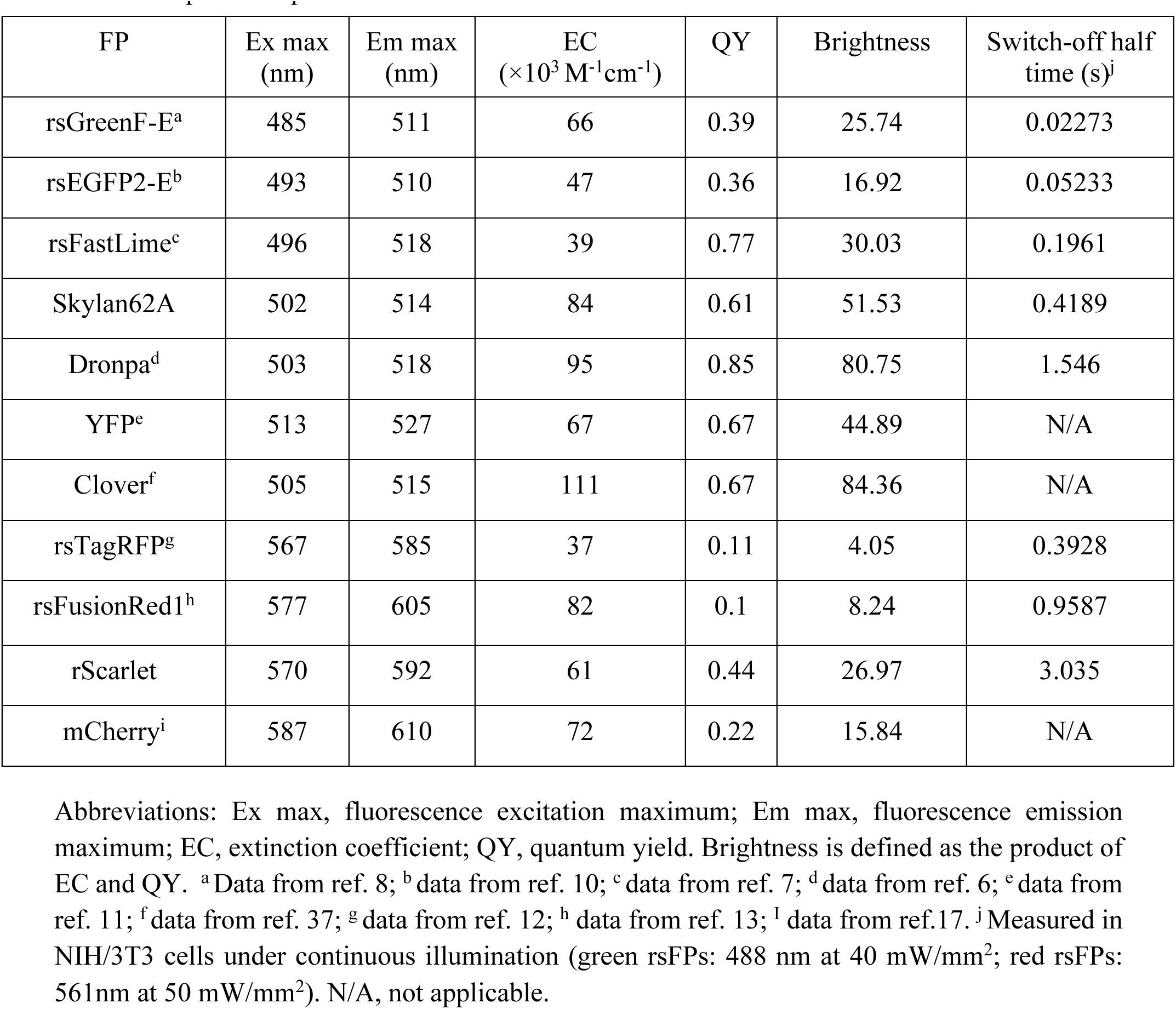
Spectroscopic characteristics of FPs used for TMI in this work.

**Table S2.**
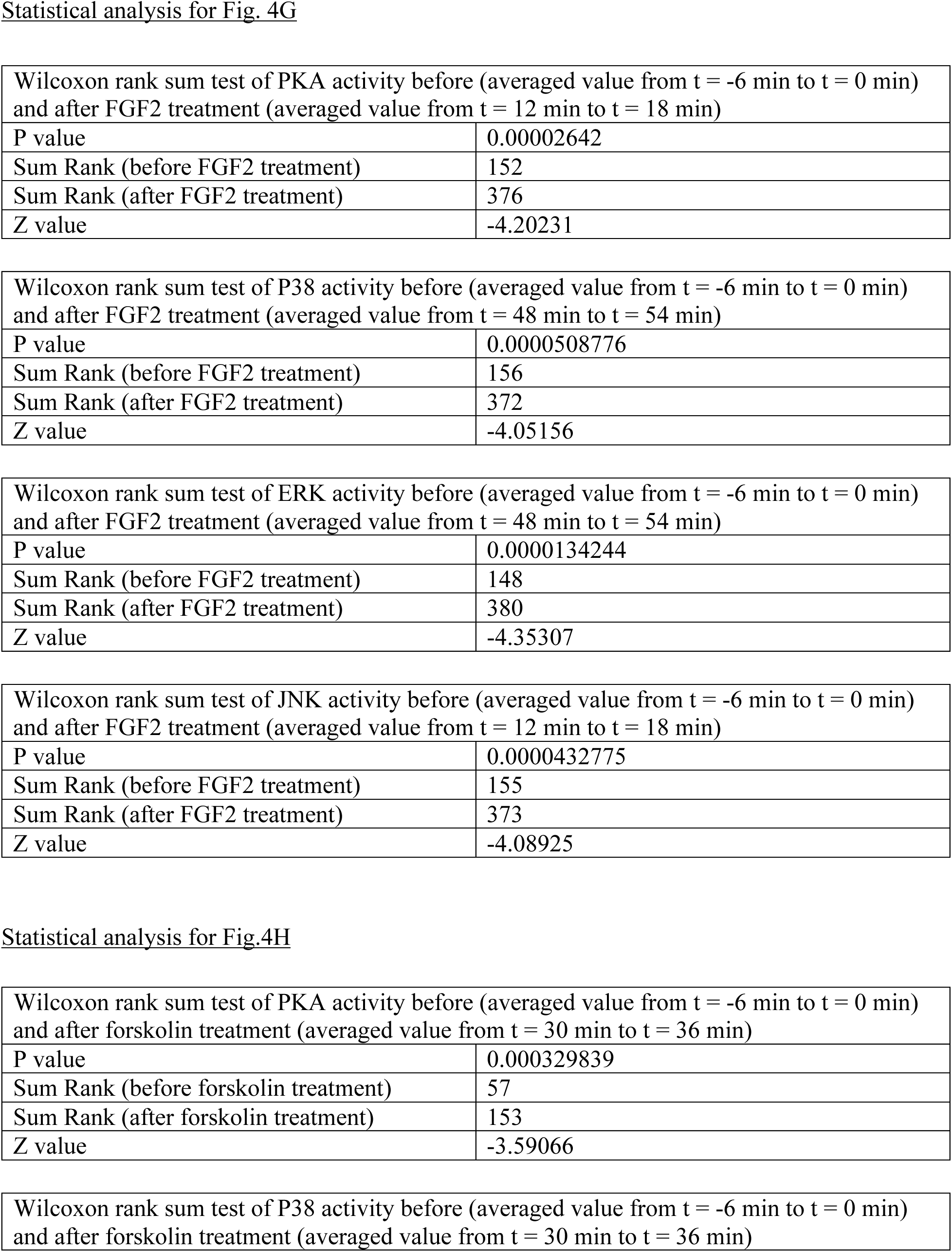
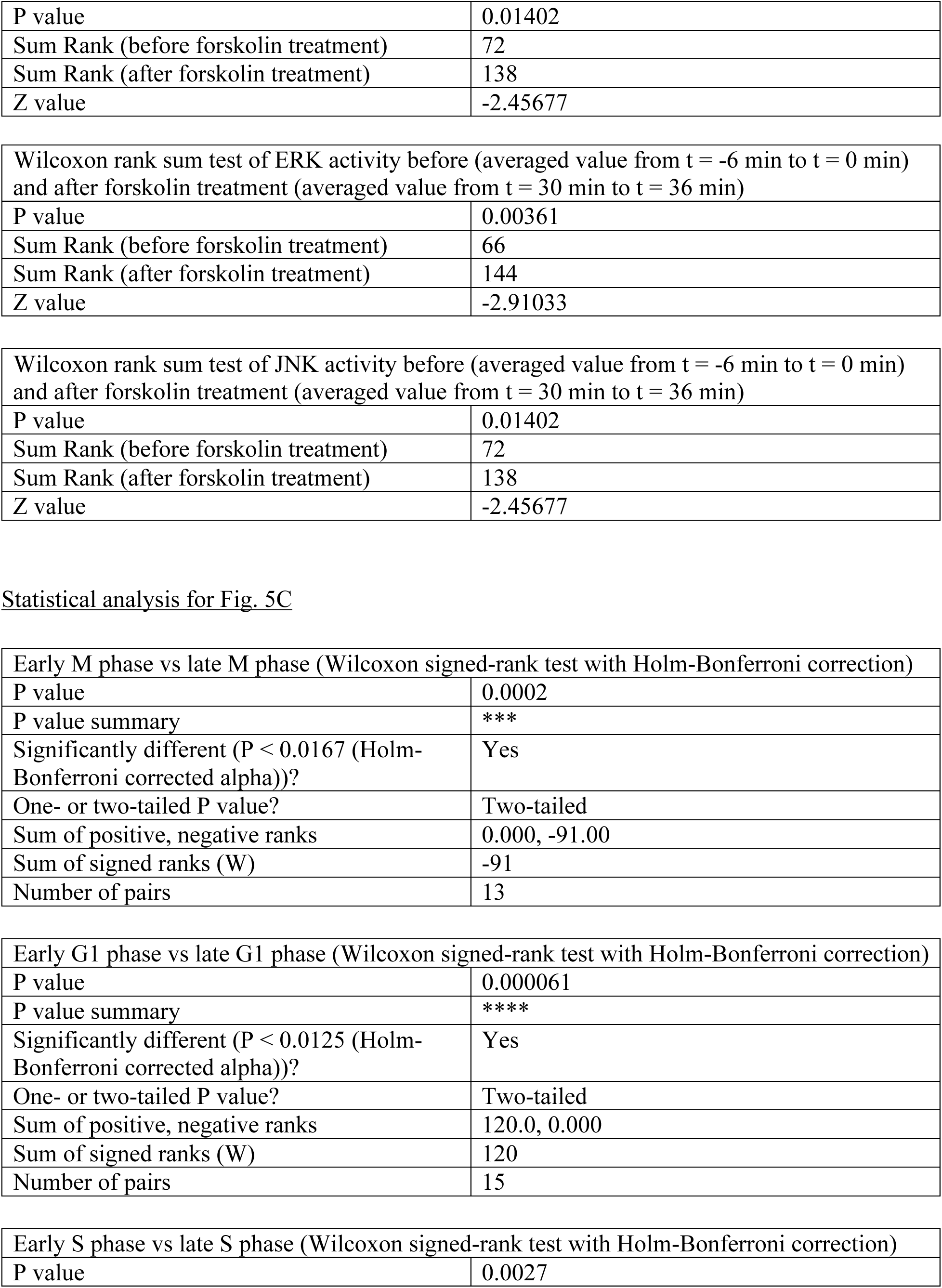
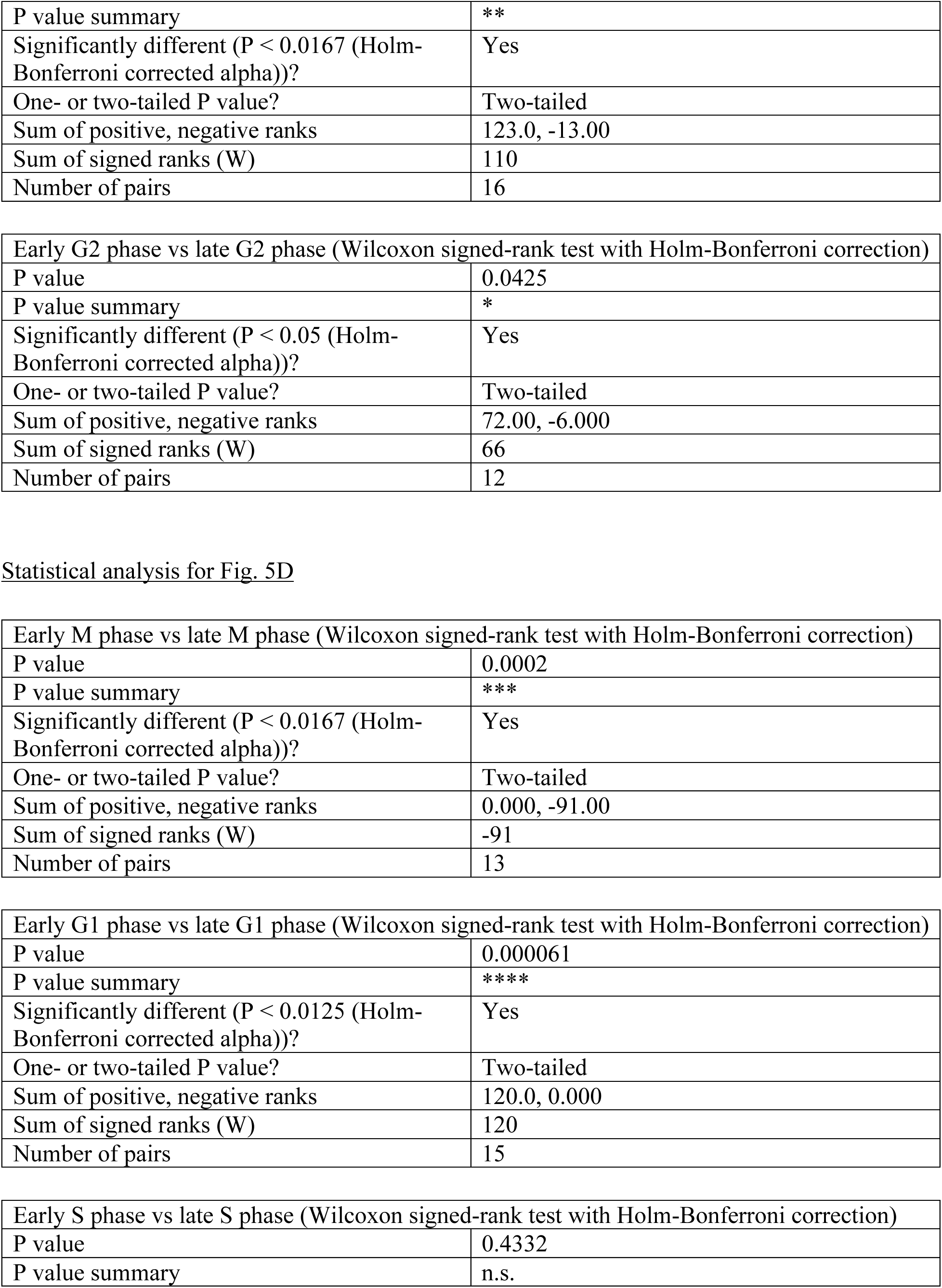
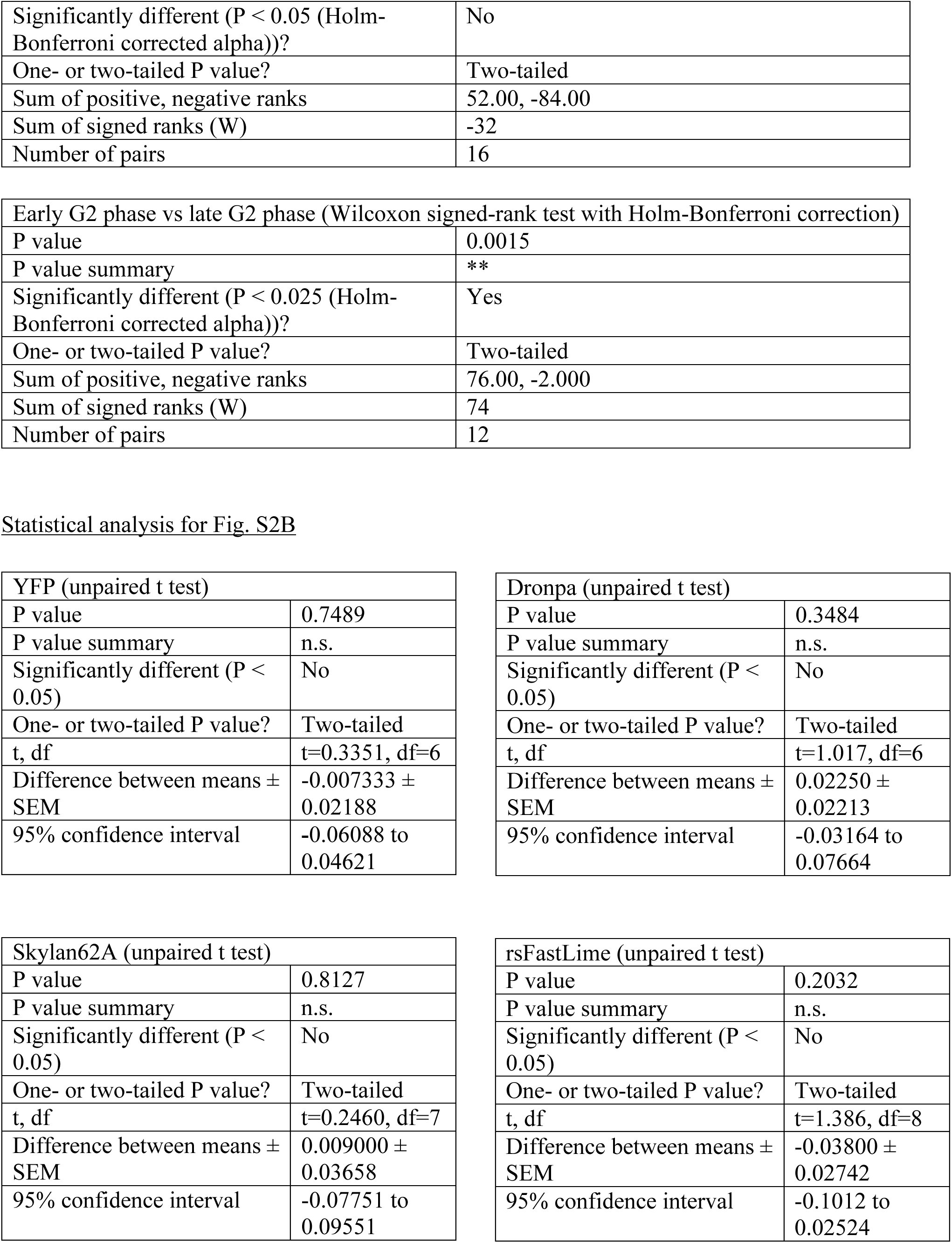
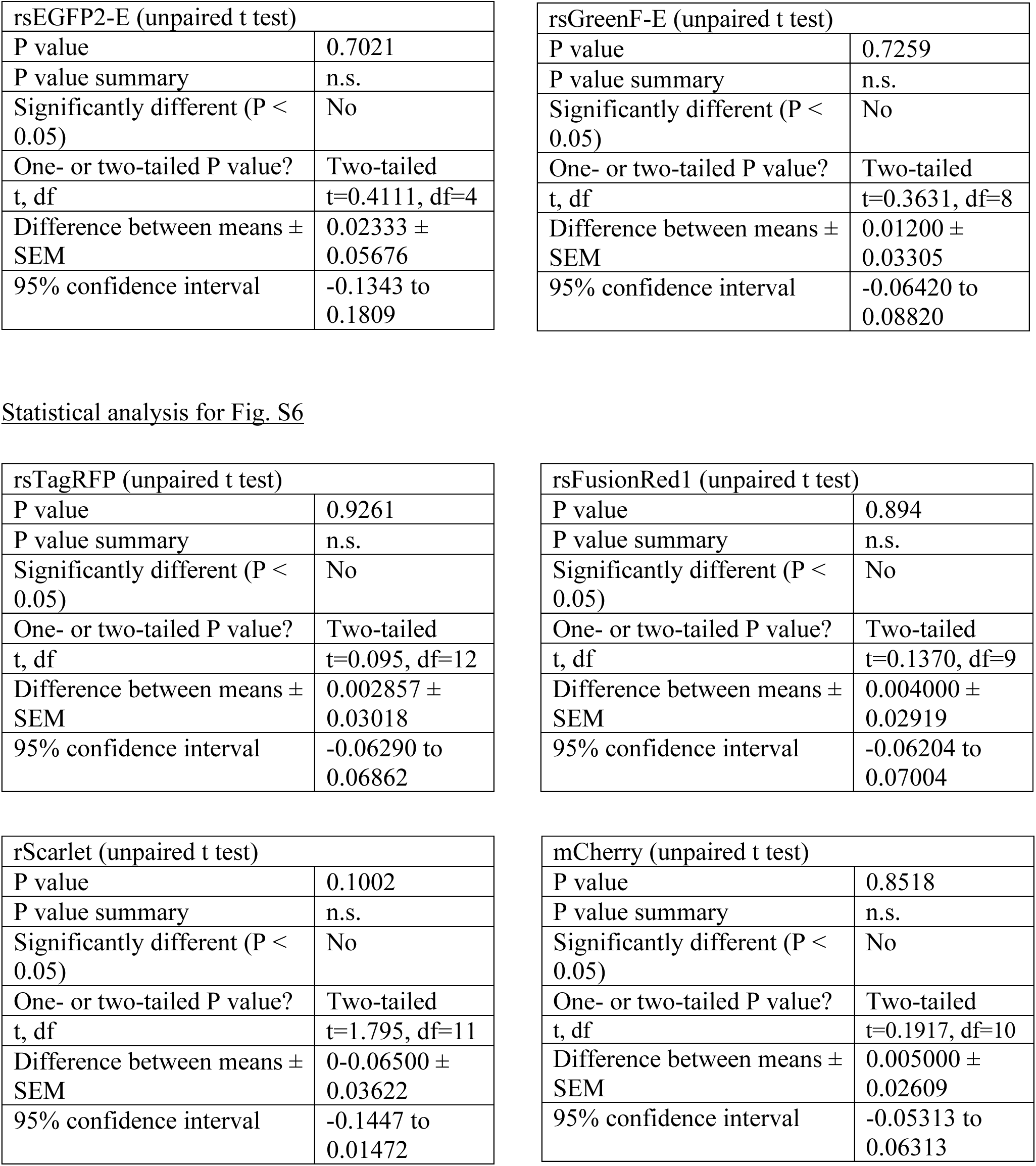
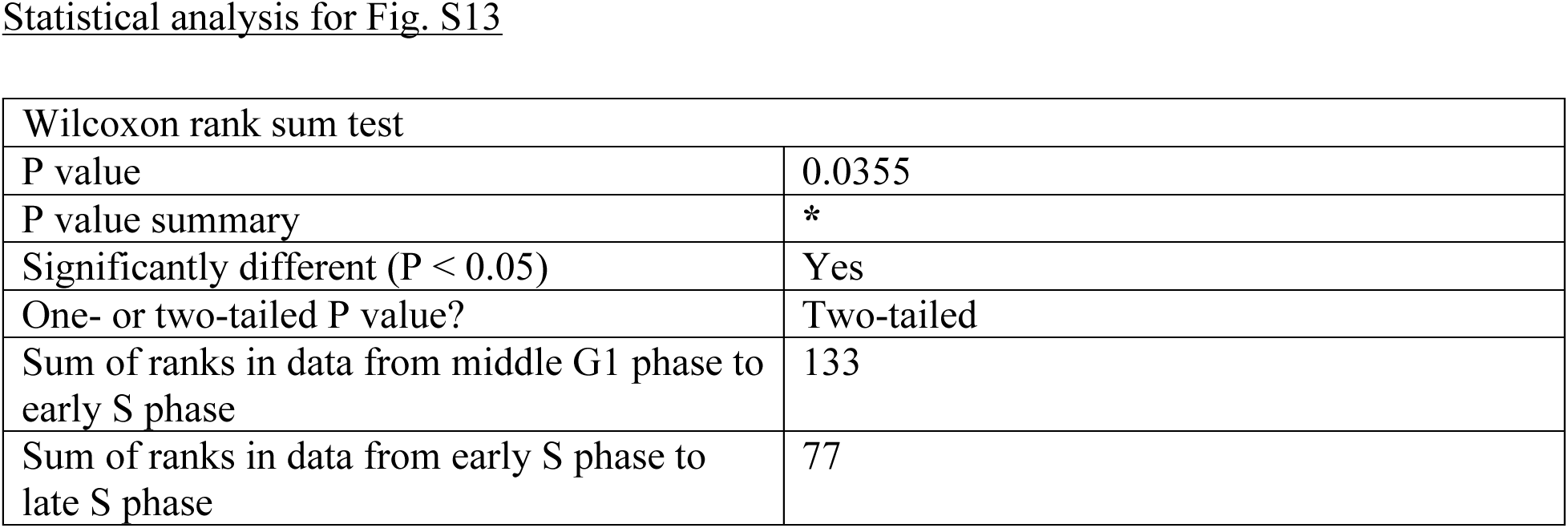
Statistical analysis.

analysis for Fig. 4G
|  |  |
| --- | --- |
| Wilcoxon rank sum test of PKA activity before (averaged value from t = -6 min to t = 0 min) and after FGF2 treatment (averaged value from t = 12 min to t = 18 min) |  |
| P value | 0.00002642 |
| Sum Rank (before FGF2 treatment) | 152 |
| Sum Rank (after FGF2 treatment) | 376 |
| Z value | -4.20231 |
| Wilcoxon rank sum test of P38 activity before (averaged value from t = -6 min to t = 0 min) and after FGF2 treatment (averaged value from t = 48 min to t = 54 min) |  |
| P value | 0.0000508776 |
| Sum Rank (before FGF2 treatment) | 156 |
| Sum Rank (after FGF2 treatment) | 372 |
| Z value | -4.05156 |
| Wilcoxon rank sum test of ERK activity before (averaged value from t = -6 min to t = 0 min) and after FGF2 treatment (averaged value from t = 48 min to t = 54 min) |  |
| P value | 0.0000134244 |
| Sum Rank (before FGF2 treatment) | 148 |
| Sum Rank (after FGF2 treatment) | 380 |
| Z value | -4.35307 |
| Wilcoxon rank sum test of JNK activity before (averaged value from t = -6 min to t = 0 min) and after FGF2 treatment (averaged value from t = 12 min to t = 18 min) |  |
| P value | 0.0000432775 |
| Sum Rank (before FGF2 treatment) | 155 |
| Sum Rank (after FGF2 treatment) | 373 |
| Z value | -4.08925 |
| Statistical analysis for Fig.4H |  |
| Wilcoxon rank sum test of PKA activity before (averaged value from t = -6 min to t = 0 min) and after forskolin treatment (averaged value from t = 30 min to t = 36 min) |  |
| P value | 0.000329839 |
| Sum Rank (before forskolin treatment) | 57 |
| Sum Rank (after forskolin treatment) | 153 |
| Z value | -3.59066 |

|  |
| --- |
| Wilcoxon rank sum test of P38 activity before (averaged value from t = -6 min to t = 0 min) and after forskolin treatment (averaged value from t = 30 min to t = 36 min) |
| P value | 0.01402 |
| Sum Rank (before forskolin treatment) | 72 |
| Sum Rank (after forskolin treatment) | 138 |
| Z value | -2.45677 |
| Wilcoxon rank sum test of ERK activity before (averaged value from t = -6 min to t = 0 min) and after forskolin treatment (averaged value from t = 30 min to t = 36 min) |  |
| P value | 0.00361 |
| Sum Rank (before forskolin treatment) | 66 |
| Sum Rank (after forskolin treatment) | 144 |
| Z value | -2.91033 |
| Wilcoxon rank sum test of JNK activity before (averaged value from t = -6 min to t = 0 min) and after forskolin treatment (averaged value from t = 30 min to t = 36 min) |  |
| P value | 0.01402 |
| Sum Rank (before forskolin treatment) | 72 |
| Sum Rank (after forskolin treatment) | 138 |
| Z value | -2.45677 |
| Statistical analysis for Fig. 5C |  |
| Early M phase vs late M phase (Wilcoxon signed-rank test with Holm-Bonferroni correction) |  |
| P value | 0.0002 |
| P value summary | *** |
| Significantly different ($P < 0.0167$ (Holm-Bonferroni corrected alpha))? | Yes |
| One- or two-tailed P value? | Two-tailed |
| Sum of positive, negative ranks | 0.000, -91.00 |
| Sum of signed ranks (W) | -91 |
| Number of pairs | 13 |
| Early G1 phase vs late G1 phase (Wilcoxon signed-rank test with Holm-Bonferroni correction) |  |
| P value | 0.000061 |
| P value summary | **** |
| Significantly different ($P < 0.0125$ (Holm-Bonferroni corrected alpha))? | Yes |
| One- or two-tailed P value? | Two-tailed |
| Sum of positive, negative ranks | 120.0, 0.000 |
| Sum of signed ranks (W) | 120 |
| Number of pairs | 15 |
| Early S phase vs late S phase (Wilcoxon signed-rank test with Holm-Bonferroni correction) |  |
| P value | 0.0027 |
| P value summary | ** |
| Significantly different (P < 0.0167 (Holm-Bonferroni corrected alpha))? | Yes |
| One- or two-tailed P value? | Two-tailed |
| Sum of positive, negative ranks | 123.0, -13.00 |
| Sum of signed ranks (W) | 110 |
| Number of pairs | 16 |
| Early G2 phase vs late G2 phase (Wilcoxon signed-rank test with Holm-Bonferroni correction) |  |
| P value | 0.0425 |
| P value summary | * |
| Significantly different (P < 0.05 (Holm-Bonferroni corrected alpha))? | Yes |
| One- or two-tailed P value? | Two-tailed |
| Sum of positive, negative ranks | 72.00, -6.000 |
| Sum of signed ranks (W) | 66 |
| Number of pairs | 12 |
| Statistical analysis for Fig. 5D |  |
| Early M phase vs late M phase (Wilcoxon signed-rank test with Holm-Bonferroni correction) |  |
| P value | 0.0002 |
| P value summary | *** |
| Significantly different (P < 0.0167 (Holm-Bonferroni corrected alpha))? | Yes |
| One- or two-tailed P value? | Two-tailed |
| Sum of positive, negative ranks | 0.000, -91.00 |
| Sum of signed ranks (W) | -91 |
| Number of pairs | 13 |
| Early G1 phase vs late G1 phase (Wilcoxon signed-rank test with Holm-Bonferroni correction) |  |
| P value | 0.000061 |
| P value summary | **** |
| Significantly different (P < 0.0125 (Holm-Bonferroni corrected alpha))? | Yes |
| One- or two-tailed P value? | Two-tailed |
| Sum of positive, negative ranks | 120.0, 0.000 |
| Sum of signed ranks (W) | 120 |
| Number of pairs | 15 |
| Early S phase vs late S phase (Wilcoxon signed-rank test with Holm-Bonferroni correction) |  |
| P value | 0.4332 |
| P value summary | n.s. |
| Significantly different ($P < 0.05$ (Holm-Bonferroni corrected alpha))? | No |
| One- or two-tailed P value? | Two-tailed |
| Sum of positive, negative ranks | 52.00, -84.00 |
| Sum of signed ranks (W) | -32 |
| Number of pairs | 16 |
| Early G2 phase vs late G2 phase (Wilcoxon signed-rank test with Holm-Bonferroni correction) |  |
| P value | 0.0015 |
| P value summary | ** |
| Significantly different ($P < 0.025$ (Holm-Bonferroni corrected alpha))? | Yes |
| One- or two-tailed P value? | Two-tailed |
| Sum of positive, negative ranks | 76.00, -2.000 |
| Sum of signed ranks (W) | 74 |
| Number of pairs | 12 |
| Statistical analysis for Fig. S2B |  |
| YFP (unpaired t test) |  |
| P value | 0.7489 |
| P value summary | n.s. |
| Significantly different ($P < 0.05$) | No |
| One- or two-tailed P value? | Two-tailed |
| t, df | t=0.3351, df=6 |
| Difference between means ± SEM | -0.007333 ± 0.02188 |
| 95% confidence interval | -0.06088 to 0.04621 |
| Dronpa (unpaired t test) |  |
| P value | 0.3484 |
| P value summary | n.s. |
| Significantly different ($P < 0.05$) | No |
| One- or two-tailed P value? | Two-tailed |
| t, df | t=1.017, df=6 |
| Difference between means ± SEM | 0.02250 ± 0.02213 |
| 95% confidence interval | -0.03164 to 0.07664 |
| Skylan62A (unpaired t test) |  |
| P value | 0.8127 |
| P value summary | n.s. |
| Significantly different ($P < 0.05$) | No |
| One- or two-tailed P value? | Two-tailed |
| t, df | t=0.2460, df=7 |
| Difference between means ± SEM | 0.009000 ± 0.03658 |
| 95% confidence interval | -0.07751 to 0.09551 |
| rsFastLime (unpaired t test) |  |
| P value | 0.2032 |
| P value summary | n.s. |
| Significantly different ($P < 0.05$) | No |
| One- or two-tailed P value? | Two-tailed |
| t, df | t=1.386, df=8 |
| Difference between means ± SEM | -0.03800 ± 0.02742 |
| 95% confidence interval | -0.1012 to 0.02524 |

| rsEGFP2-E (unpaired t test) |  |
| --- | --- |
| P value | 0.7021 |
| P value summary | n.s. |
| Significantly different (P < 0.05) | No |
| One- or two-tailed P value? | Two-tailed |
| t, df | t=0.4111, df=4 |
| Difference between means ± SEM | 0.02333 ± 0.05676 |
| 95% confidence interval | -0.1343 to 0.1809 |

| rsGreenF-E (unpaired t test) |  |
| --- | --- |
| P value | 0.7259 |
| P value summary | n.s. |
| Significantly different (P < 0.05) | No |
| One- or two-tailed P value? | Two-tailed |
| t, df | t=0.3631, df=8 |
| Difference between means ± SEM | 0.01200 ± 0.03305 |
| 95% confidence interval | -0.06420 to 0.08820 |

Statistical analysis for Fig. S6
| rsTagRFP (unpaired t test) |  |
| --- | --- |
| P value | 0.9261 |
| P value summary | n.s. |
| Significantly different (P < 0.05) | No |
| One- or two-tailed P value? | Two-tailed |
| t, df | t=0.095, df=12 |
| Difference between means ± SEM | 0.002857 ± 0.03018 |
| 95% confidence interval | -0.06290 to 0.06862 |

| rsFusionRed1 (unpaired t test) |  |
| --- | --- |
| P value | 0.894 |
| P value summary | n.s. |
| Significantly different (P < 0.05) | No |
| One- or two-tailed P value? | Two-tailed |
| t, df | t=0.1370, df=9 |
| Difference between means ± SEM | 0.004000 ± 0.02919 |
| 95% confidence interval | -0.06204 to 0.07004 |

| rScarlet (unpaired t test) |  |
| --- | --- |
| P value | 0.1002 |
| P value summary | n.s. |
| Significantly different (P < 0.05) | No |
| One- or two-tailed P value? | Two-tailed |
| t, df | t=1.795, df=11 |
| Difference between means ± SEM | 0-0.06500 ± 0.03622 |
| 95% confidence interval | -0.1447 to 0.01472 |

| mCherry (unpaired t test) |  |
| --- | --- |
| P value | 0.8518 |
| P value summary | n.s. |
| Significantly different (P < 0.05) | No |
| One- or two-tailed P value? | Two-tailed |
| t, df | t=0.1917, df=10 |
| Difference between means ± SEM | 0.005000 ± 0.02609 |
| 95% confidence interval | -0.05313 to 0.06313 |
| Statistical analysis for Fig. S13 |  |
| Wilcoxon rank sum test |  |
| P value | 0.0355 |
| P value summary | * |
| Significantly different ($P < 0.05$) | Yes |
| One- or two-tailed P value? | Two-tailed |
| Sum of ranks in data from middle G1 phase to early S phase | 133 |
| Sum of ranks in data from early S phase to late S phase | 77 |

## References

1. C. Linghu, S. L. Johnson, P. A. Valdes, O. A. Shemesh, W. M. Park, D. Park, K. D. Piatkevich, A. T. Wassie, Y. Liu, B. An, S. A. Barnes, O. T. Celiker, C.-C. Yao, C.-C. J. Yu, R. Wang, K. P. Adamala, M. F. Bear, A. E. Keating, E. S. Boyden, Spatial Multiplexing of Fluorescent Reporters for Imaging Signaling Network Dynamics. Cell. 183,1682–1698.e24 (2020).

2. S. Mehta, Y. Zhang, R. H. Roth, J.-F. Zhang, A. Mo, B. Tenner, R. L. Huganir, J. Zhang, Single-fluorophore biosensors for sensitive and multiplexed detection of signalling activities. Nat. Cell Biol. 20, 1215–1225 (2018).

3. K. Chen, R. Yan, L. Xiang, K. Xu, Excitation spectral microscopy for highly multiplexed fluorescence imaging and quantitative biosensing. Light Sci Appl. 10, 97 (2021).

4. D. M. Shcherbakova, O. V. Stepanenko, K. K. Turoverov, V. V. Verkhusha, Near-Infrared Fluorescent Proteins: Multiplexing and Optogenetics across Scales. Trends Biotechnol. 36, 1230–1243 (2018).

5. F. Chen, P. W. Tillberg, E. S. Boyden, Optical imaging. Expansion microscopy. Science. 347, 543–548 (2015).

6. R. Ando, H. Mizuno, A. Miyawaki, Regulated fast nucleocytoplasmic shuttling observed by reversible protein highlighting. Science. 306, 1370–1373 (2004).

7. A. C. Stiel, S. Trowitzsch, G. Weber, M. Andresen, C. Eggeling, S. W. Hell, S. Jakobs, M. C. Wahl, 1.8 A bright-state structure of the reversibly switchable fluorescent protein Dronpa guides the generation of fast switching variants. Biochem. J. 402, 35–42 (2007).

8. T. Roebroek, S. Duwé, W. Vandenberg, P. Dedecker, Reduced Fluorescent Protein Switching Fatigue by Binding-Induced Emissive State Stabilization. Int. J. Mol. Sci. 18 (2017).

9. X. Zhang, M. Zhang, D. Li, W. He, J. Peng, E. Betzig, P. Xu, Highly photostable, reversibly photoswitchable fluorescent protein with high contrast ratio for live-cell superresolution microscopy. Proc. Natl. Acad. Sci. U. S. A. 113, 10364–10369 (2016).

10. T. Grotjohann, I. Testa, M. Reuss, T. Brakemann, C. Eggeling, S. W. Hell, S. Jakobs, rsEGFP2 enables fast RESOLFT nanoscopy of living cells. Elife. 1, e00248 (2012).

11. M. Ormö, A. B. Cubitt, K. Kallio, L. A. Gross, R. Y. Tsien, S. J. Remington, Crystal structure of the Aequorea victoria green fluorescent protein. Science. 273, 1392–1395 (1996).

12. S. Pletnev, F. V. Subach, Z. Dauter, A. Wlodawer, V. V. Verkhusha, A structural basis for reversible photoswitching of absorbance spectra in red fluorescent protein rsTagRFP. J. Mol. Biol. 417, 144–151 (2012).

13. F. Pennacchietti, E. O. Serebrovskaya, A. R. Faro, I. I. Shemyakina, N. G. Bozhanova, A. A. Kotlobay, N. G. Gurskaya, A. Bodén, J. Dreier, D. M. Chudakov, K. A. Lukyanov, V. V. Verkhusha, A. S. Mishin, I. Testa, Fast reversibly photoswitching red fluorescent proteins for live-cell RESOLFT nanoscopy. Nat. Methods. 15, 601–604 (2018).

14. D. S. Bindels, L. Haarbosch, L. van Weeren, M. Postma, K. E. Wiese, M. Mastop, S. Aumonier, G. Gotthard, A. Royant, M. A. Hink, T. W. J. Gadella Jr, mScarlet: a bright monomeric red fluorescent protein for cellular imaging. Nat. Methods. 14, 53–56 (2017).

15. A. C. Stiel, M. Andresen, H. Bock, M. Hilbert, J. Schilde, A. Schönle, C. Eggeling, A. Egner, S. W. Hell, S. Jakobs, Generation of monomeric reversibly switchable red fluorescent proteins for far-field fluorescence nanoscopy. Biophys. J. 95, 2989–2997 (2008).

16. F. Lavoie-Cardinal, N. A. Jensen, V. Westphal, A. C. Stiel, A. Chmyrov, J. Bierwagen, I. Testa, S. Jakobs, S. W. Hell, Two-color RESOLFT nanoscopy with green and red fluorescent photochromic proteins. Chemphyschem. 15, 655–663 (2014).

17. N. C. Shaner, R. E. Campbell, P. A. Steinbach, B. N. G. Giepmans, A. E. Palmer, R. Y. Tsien, Improved monomeric red, orange and yellow fluorescent proteins derived from aaDiscosoma sp. red fluorescent protein. Nat. Biotechnol. 22, 1567–1572 (2004).

18. J. Livet, T. A. Weissman, H. Kang, R. W. Draft, J. Lu, R. A. Bennis, J. R. Sanes, J. W. Lichtman, Transgenic strategies for combinatorial expression of fluorescent proteins in the nervous system. Nature. 450, 56–62 (2007).

19. R. W. Köster, S. E. Fraser, Tracing transgene expression in living zebrafish embryos. Dev. Biol. 233, 329–346 (2001).

20. I. Formella, A. J. Svahn, R. A. W. Radford, E. K. Don, N. J. Cole, A. Hogan, A. Lee, R. S. Chung, M. Morsch, Real-time visualization of oxidative stress-mediated neurodegeneration of individual spinal motor neurons in vivo. Redox Biol. 19, 226–234 (2018).

21. B. T. Bajar, A. J. Lam, R. K. Badiee, Y.-H. Oh, J. Chu, X. X. Zhou, N. Kim, B. B. Kim, M. Chung, A. L. Yablonovitch, B. F. Cruz, K. Kulalert, J. J. Tao, T. Meyer, X.-D. Su, M. Z. Lin, Fluorescent indicators for simultaneous reporting of all four cell cycle phases. Nat. Methods. 13, 993–996 (2016).

22. S. Regot, J. J. Hughey, B. T. Bajar, S. Carrasco, M. W. Covert, High-sensitivity measurements of multiple kinase activities in live single cells. Cell. 157, 1724–1734 (2014).

23. J. P. Pursiheimo, M. Jalkanen, K. Taskén, P. Jaakkola, Involvement of protein kinase A in fibroblast growth factor-2-activated transcription. Proc. Natl. Acad. Sci. U. S. A. 97, 168–173 (2000).

24. M. P. Lichtenstein, J. L. M. Madrigal, A. Pujol, E. Galea, JNK/ERK/FAK mediate promigratory actions of basic fibroblast growth factor in astrocytes via CCL2 and COX2. Neurosignals. 20, 86–102 (2012).

25. B. S. Kim, J.-Y. Park, H.-J. Kang, H.-J. Kim, J. Lee, Fucoidan/FGF-2 induces angiogenesis through JNK- and p38-mediated activation of AKT/MMP-2 signalling. Biochem. Biophys. Res. Commun. 450, 1333–1338 (2014).

26. S. Kanazawa, T. Fujiwara, S. Matsuzaki, K. Shingaki, M. Taniguchi, S. Miyata, M. Tohyama, Y. Sakai, K. Yano, K. Hosokawa, T. Kubo, bFGF regulates PI3-kinase-Rac1-JNK pathway and promotes fibroblast migration in wound healing. PLoS One. 5, e12228 (2010).

27. K. H. Park, H. J. Park, K. S. Shin, H. S. Choi, M. Kai, M. K. Lee, Modulation of PC12 cell viability by forskolin-induced cyclic AMP levels through ERK and JNK pathways: an implication for L-DOPA-induced cytotoxicity in nigrostriatal dopamine neurons. Toxicol. Sci. 128, 247–257 (2012).

28. M. P. Delghandi, M. Johannessen, U. Moens, The cAMP signalling pathway activates CREB through PKA, p38 and MSK1 in NIH 3T3 cells. Cell. Signal. 17, 1343–1351 (2005).

29. S. L. Spencer, S. D. Cappell, F.-C. Tsai, K. W. Overton, C. L. Wang, T. Meyer, The proliferation-quiescence decision is controlled by a bifurcation in CDK2 activity at mitotic exit. Cell. 155, 369–383 (2013).

30. H. W. Yang, S. D. Cappell, A. Jaimovich, C. Liu, M. Chung, L. H. Daigh, L. R. Pack, Y. Fan, S. Regot, M. Covert, T. Meyer, Stress-mediated exit to quiescence restricted by increasing persistence in CDK4/6 activation. Elife. 9 (2020).

31. B. Stern, P. Nurse, A quantitative model for the cdc2 control of S phase and mitosis in fission yeast. Trends Genet. 12, 345–350 (1996).

32. D. Coudreuse, P. Nurse, Driving the cell cycle with a minimal CDK control network. Nature. 468, 1074–1079 (2010).

33. Y. Qian, D. M. O. Cosio, K. D. Piatkevich, S. Aufmkolk, W.-C. Su, O. T. Celiker, A. Schohl, M. H. Murdock, A. Aggarwal, Y.-F. Chang, P. W. Wiseman, E. S. Ruthazer, E. S. Boyden, R. E. Campbell, Improved genetically encoded near-infrared fluorescent calcium ion indicators for in vivo imaging. PLoS Biol. 18, e3000965 (2020).

34. K. Harada, M. Ito, X. Wang, M. Tanaka, D. Wongso, A. Konno, H. Hirai, H. Hirase, T. Tsuboi, T. Kitaguchi, Red fluorescent protein-based cAMP indicator applicable to optogenetics and in vivo imaging. Sci. Rep. 7, 7351 (2017).

35. G. Baillie, S. J. MacKenzie, M. D. Houslay, Phorbol 12-myristate 13-acetate triggers the protein kinase A-mediated phosphorylation and activation of the PDE4D5 cAMP phosphodiesterase in human aortic smooth muscle cells through a route involving extracellular signal regulated kinase (ERK). Mol. Pharmacol. 60, 1100–1111 (2001).

36. V. Sriraman, S. R. Modi, Y. Bodenburg, L. A. Denner, R. J. Urban, Identification of ERK and JNK as signaling mediators on protein kinase C activation in cultured granulosa cells. Mol. Cell. Endocrinol. 294, 52–60 (2008).

## Supplementary References

37. A. J. Lam, F. St-Pierre, Y. Gong, J. D. Marshall, P. J. Cranfill, M. A. Baird, M. R. McKeown, J. Wiedenmann, M. W. Davidson, M. J. Schnitzer, R. Y. Tsien, M. Z. Lin, Improving FRET dynamic range with bright green and red fluorescent proteins. Nat. Methods. 9, 1005–1012 (2012).

38. N. C. Shaner, M. Z. Lin, M. R. McKeown, P. A. Steinbach, K. L. Hazelwood, M. W. Davidson, R. Y. Tsien, Improving the photostability of bright monomeric orange and red fluorescent proteins. Nat. Methods. 5, 545–551 (2008).

39. D. Shcherbo, C. S. Murphy, G. V. Ermakova, E. A. Solovieva, T. V. Chepurnykh, A. S. Shcheglov, V. V. Verkhusha, V. Z. Pletnev, K. L. Hazelwood, P. M. Roche, S. Lukyanov, A. G. Zaraisky, M. W. Davidson, D. M. Chudakov, Far-red fluorescent tags for protein imaging in living tissues. Biochem. J. 418, 567–574 (2009).

40. E. M. Merzlyak, J. Goedhart, D. Shcherbo, M. E. Bulina, A. S. Shcheglov, A. F. Fradkov, A. Gaintzeva, K. A. Lukyanov, S. Lukyanov, T. W. J. Gadella, D. M. Chudakov, Bright monomeric red fluorescent protein with an extended fluorescence lifetime. Nat. Methods. 4, 555–557 (2007).

41. G. C. H. Mo, C. Posner, E. A. Rodriguez, T. Sun, J. Zhang, A rationally enhanced red fluorescent protein expands the utility of FRET biosensors. Nat. Commun. 11, 1848 (2020).

42. I. I. Shemiakina, G. V. Ermakova, P. J. Cranfill, M. A. Baird, R. A. Evans, E. A. Souslova, D. B. Staroverov, A. Y. Gorokhovatsky, E. V. Putintseva, T. V. Gorodnicheva, T. V. Chepurnykh, L. Strukova, S. Lukyanov, A. G. Zaraisky, M. W. Davidson, D. M. Chudakov, D. Shcherbo, A monomeric red fluorescent protein with low cytotoxicity. Nat. Commun. 3, 1204 (2012).

43. M. Z. Lin, M. R. McKeown, H.-L. Ng, T. A. Aguilera, N. C. Shaner, R. E. Campbell, S. R. Adams, L. A. Gross, W. Ma, T. Alber, R. Y. Tsien, Autofluorescent proteins with excitation in the optical window for intravital imaging in mammals. Chem. Biol. 16, 1169–1179 (2009).

44. J. Chu, R. D. Haynes, S. Y. Corbel, P. Li, E. González-González, J. S. Burg, N. J. Ataie, A. J. Lam, P. J. Cranfill, M. A. Baird, M. W. Davidson, H.-L. Ng, K. C. Garcia, C. H. Contag, K. Shen, H. M. Blau, M. Z. Lin, Non-invasive intravital imaging of cellular differentiation with a bright red-excitable fluorescent protein. Nat. Methods. 11, 572–578 (2014).

45. L. A. Gross, G. S. Baird, R. C. Hoffman, K. K. Baldridge, R. Y. Tsien, The structure of the chromophore within DsRed, a red fluorescent protein from coral. Proc. Natl. Acad. Sci. U. S. A. 97, 11990–11995 (2000).

46. J. M. Kendall, R. L. Dormer, A. K. Campbell, Targeting aequorin to the endoplasmic reticulum of living cells. Biochem. Biophys. Res. Commun. 189, 1008–1016 (1992).

47. N. M. Sherer, M. J. Lehmann, L. F. Jimenez-Soto, A. Ingmundson, S. M. Horner, G. Cicchetti, P. G. Allen, M. Pypaert, J. M. Cunningham, W. Mothes, Visualization of retroviral replication in living cells reveals budding into multivesicular bodies. Traffic. 4, 785–801 (2003).

48. Y. Kimura, C. Satou, S.-I. Higashijima, V2a and V2b neurons are generated by the final divisions of pair-producing progenitors in the zebrafish spinal cord. Development. 135, 3001–3005 (2008).

49. K. M. Kwan, E. Fujimoto, C. Grabher, B. D. Mangum, M. E. Hardy, D. S. Campbell, J. M. Parant, H. J. Yost, J. P. Kanki, C.-B. Chien, The Tol2kit: a multisite gateway-based construction kit for Tol2 transposon transgenesis constructs. Dev. Dyn. 236, 3088–3099 (2007).

50. S. Fisher, E. A. Grice, R. M. Vinton, S. L. Bessling, A. Urasaki, K. Kawakami, A. S. McCallion, Evaluating the biological relevance of putative enhancers using Tol2 transposon-mediated transgenesis in zebrafish. Nat. Protoc. 1, 1297–1305 (2006).

51. U. Schmidt, M. Weigert, C. Broaddus, G. Myers, in Medical Image Computing and Computer Assisted Intervention – MICCAI 2018 (Springer International Publishing, 2018), pp. 265–273.

